# Simulation of Protein Structure using a Coarse-Grained Potential incorporating the Backbone Dihedral Interactions

**DOI:** 10.1101/2025.08.20.671185

**Authors:** Kanika Kole, Abhik Ghosh Moulick, Jaydeb Chakrabarti

## Abstract

Many biologically relevant processes occur on time and length scales which are far beyond the reach of atomistic simulations. These processes include large protein dynamics and the self-assembly of biological materials. Coarse-grained molecular modeling allows computer simulations on length and time scales 2–3 orders of magnitude larger than atomistic simulations, bridging the gap between the atomistic and mesoscopic scales. However, the structural information involving the dihedral angles is lost in coarse-graining. We develop a simple coarse-grained protein model with structural information in explicit solvent. We represent the center of mass of each residue as a polymer bead and water oxygen as a solvent bead. Each polymer bead has five degrees of freedom: position of the center and two additional variables for the backbone dihedral angles. All interaction parameters for bonded, non-bonded, dihedral coupling and bead-solvent interactions are derived from the equilibrated all-atom molecular dynamics simulation trajectory. We find that our coarse-grained approach reproduces residue-level structural information that closely matches the crystal structures and all-atom simulation results.

## 1 Introduction

Solution phase behaviours of bio-macromolecular systems, like aggregation of proteins, involve a large number of protein molecules. Microscopic description of such phenomenon is far beyond the reach of full microscopic all-atom (AA) models ^1^ and typically involves coarse-grained (CG) models which describe complex systems at large length and time scales ^2–8^ by reducing the number of degrees of freedom, a number of atoms being grouped together into a unit ^9,10^. However, the explicit information on atomic planes and the dihedral angles between atomic planes ^11^, the key ingredients to describe the spatial conformations of complex molecules, are lost due to grouping of atoms. Consequently, CG models have inherent limitations to adequately describe cases where the bio-molecular conformations lie at the center stage.

The backbone dihedral angles describe protein conformations. Proteins which lack a stable, well-defined three-dimensional structure, like the intrinsically disordered proteins (IDP) ^12^ undergo fluctuations among conformations even in physiological conditions ^13^. They often have a tendency to aggregate in the solution phase ^14^, leading to pathological conditions ^14–16^. Sometimes proteins, like milk protein alpha-lactalbumin in denaturing conditions, form a molten globule state in which local stable structure gets disrupted while retaining overall secondary structure. Molten globule proteins are capable of ligand binding ^14,15^ and form aggregation in solution phases. The conformation fluctuations in such proteins are extremely important to understand their solution phase properties like aggregation and their functions.

A wide variety of CG models has been introduced to capture specific aspects for bio-molecular systems, like proteins ^17^, nucleic acids ^17–19^, lipid membranes ^20,21^, carbohydrates ^22^, water ^23,24^, and so on. In highly simplified lattice protein-like hydrophobic-hydrophilic (HP) models ^25–28^, each amino acid is either represented as a solvophobic or polar entity, but ignores protein backbone information. More detailed CG protein models adopt different levels of simplified polypeptide representation ^29^. The main chain is represented by either all heavy atoms or one to two united atoms per residue, while the side chain is typically replaced by one or two united atoms ^29–31^. Realistic CG models, like the Martini ^13,32^, the SIRAH ^19,34^, the SPICA, the UNRES and so on, typically derive their parameters from detailed theoretical models, such as atomistic or quantum mechanical simulations ^1^. However, each of these approaches has inherent limitations to obtain reliable dihedral angle as well as secondary-structure information directly. For force fields such as MARTINI or SIRAH, this limitation is commonly addressed through backmapping ^35^, a procedure in which atomistic (AA) structures are reconstructed from CG snapshots or trajectories. However, this is quite challenging because back-mapping is not unique and often depends on methodological choices and structural restraints, which can introduce biases and additional sources of uncertainty.

There is a clear need for a CG force fields to describe secondary structure of proteins which retain sufficient information such as backbone dihedral angle without relying on back-mapping. Goddard III and coworkers introduce a CG Dihedral Probability Grid Monte Carlo (DPG-MC) approach to generate polypeptides ^36^ and protein structures ^37^. C*α* atoms are used to represent the protein backbone and pairwise model side-chain–side-chain interactions are employed. An internal-coordinate search algorithm is employed that is guided by residue-specific dihedral angle probability distributions derived from known protein structures in the Protein Data Bank (PDB). The initial Cartesian coordinates of the polypeptides and proteins are constructed using BIOGRAF ^38^, after which the secondary structures are randomly assigned to each residue ^36^. The net charge of the peptide or protein is determined using a Charge-Equilibration scheme. The energy of the resulting structure is then minimized to convergence using the steepest-descent method, followed by conjugate-gradient minimization. The structures are updated using the DPG-MC simulations. At the end of each DPG-MC run, the lowest-energy conformation is further refined by energy minimization using steepest-descent followed by conjugate-gradient methods until convergence. This model thus integrates protein database knowledge with conformational search methodologies. However, the available PDB data are limited, particularly for intrinsically disordered proteins (IDPs). Moreover, since solvent atoms are not treated explicitly, the interactions between the amino-acid residues and solvent cannot be treated.

With this backdrop, we build a simple CG meso-scopic polymer model for protein to capture the structural aspects utilizing statistical mechanics based physical interactions derived from fully microscopic all-atom (AA) simulations. As a first step to find a suitable model to describe conformation fluctuations in large number of protein molecules aggregating in a solution, we explore how to extract at a minimal level the structural information of a protein molecule without sacrificing the inherent advantages of a meso-scopic CG model. Fig. 1(a) shows a schematic of our model, illustrating polymer beads, connected by springs with finite stretching and bending elastic constants and water oxygen by isolated spheres. The key features of our model are as follows: (1) Each polymer bead represents the centre of mass of a protein residue as in the primary sequence of the protein and the size is given by the radius of gyration of the residue consider only the heavy atoms. We assign five variables to each bead: three for the coordinate of the center of mass (COM) and two degrees of freedom corresponding to the main chain backbone dihedral angles (*ϕ* and *ψ*). (2) The beads are classified based on solvent interactions: The solvophobic beads repel water oxygen which correspond to the hydrophobic residues, while the sovophilic beads attract water oxygen, corresponding to the hydrophilic residues in a given solvent condition. (3) For a given coarse-grained variable Γ in the model, we compute the distribution of the variable *H*(Γ) over equilibrated conformations of the protein in the AA simulations. The equilibrium value and energy associated with Γ are calculated using the Boltzmann formula *F*(Γ) = −*k*_*B*_*TlnH*(Γ). The bead-solvent interactions are calculated from the distribution of water molecules around the COM of a residue over equilibrium AA trajectory. The dihedral interactions are computed from the joint probability distributions of different dihedral angles over AA trajectory and grouped together depending on solvophobic and solvophilic property of the residues. All other relevant CG model parameters, like the bead size, the backbone elastic constants, bead-bead non-bonded interactions and solvent-solvent interactions are computed using distributions of the relevant variables.

**Fig. 1.**
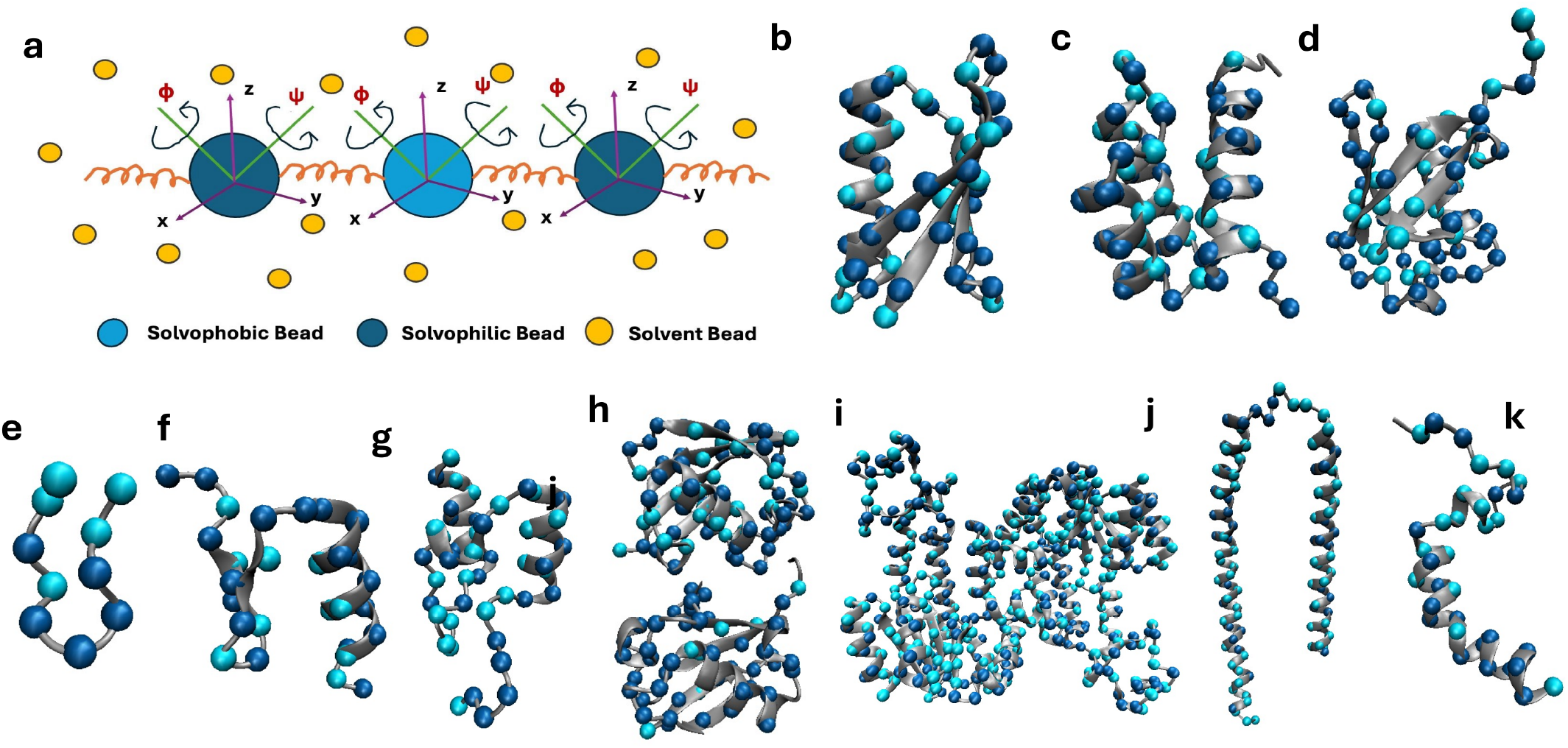
(a)A schematic illustrating the main-chain dihedral angles *ϕ* and *ψ* associated with each polymer bead. Crystal structure of (b) GB3 (PDB ID:2OED), (c) homeodomain (PDB ID: 1K61), (d) ubiquitin (PDB ID: 1UBQ), (e) Chignolin protein (PDB ID:5AWL), (f) BBL protein (PDB ID: 2WXC), (g) protein BBA(PDB ID: 1FME), (h) di-ubiquitin (PDB ID: 7S6O) (i)adenylate kinase (PDB ID:4AKE), (j) *α*-synuclein (PDB ID: 1XQ8), (k) *λ* N (PDB ID:1QFQ) protein. Coarse-grained (CG) beads representing solvophobic and solvophilic residues are depicted in van der Waals (vdW) representation.

We then perform MC simulations of the model polymer based on Metropolis sampling to generate equilibrium structure of the chain ^39,40^ in which Cartesian coordinates and backbone dihedral angles are updated at each MC step to generate conformations as per the energy costs in the CG model interactions. We test our method on GB3, a small protein with 56 amino acid residues, widely used as a model structure ^41^ in CG simulations due to its small size, well-defined structure and well-documented folding mechanism. We calculate CG model parameters from the AA simulations data of GB3. We then perform CG MC simulations using the CG parameters determined from the AA trajectory of GB3 and find that the structural elements of the proteins are well captured when compared to their crystal structure and AA data. We further apply our method using CG parameters derived from the GB3 all-atom (AA) trajectory to several well-structured proteins, like Homeodomain, Ubiquitin, Chignolin, B domain of Protein L (BBL) and Beta-Beta-Alpha (BBA). We find that the structural data for two-domain proteins, like K48-linked Di-ubiquitin and Adenylate kinase can also be captured using CG parameters derived from the GB3 force field. We further study a couple of IDPs, *α*-synuclein (*α*S) and *λ* N. AA simulations of *α*S yield the elastic constants which are smaller compared to well-folded proteins reflecting their high flexibilities. The CG simulations with these parameters show primarily unfolded structures in both *α*S and *λ* N in agreement to the AA data.

## 2 Methods

### 2.1 System preparation

We study the following systems: (1) GB3 (PDB ID: 2OED) ^42^, a small globular protein consisting of 56 residues, (2) homeodomain protein, another small globular protein containing 58 residues, the initial structure of homeodomain protein (*α*2D) is taken from the PDB structure 1K61^43^ without DNA and another homeodomain protein (*α*2B) and (3) ubiquitin (PDB ID: 1UBQ) ^44^, a small structural protein containing 76 residues. We further consider small folded proteins, (4) Chignolin protein (PDB ID:5AWL) ^45^, (5) the BBL protein (PDB ID: 2WXC ^46^) and (6) protein BBA, which contains a mixed *α*/*β* structure (PDB ID: 1FME) ^47^. Owing to their well-characterized structures and dynamics, these proteins have been widely used to explore protein folding and unfolding mechanisms in the fields of chemistry and biochemistry. Then, we apply our model on multi-domain proteins, (7) di-ubiquitin (PDB ID: 7S6O, linked via K48) ^48^, a multidomain protein consisting of 148 residues and (8) adenylate kinase, a multidomain protein of 428 residues from the kinase family (PDB ID: 4AKE) ^49^ The crystal structures corresponding to all proteins are shown in Figs. 1(b–k). We study two IDPs, (9) *α*-synuclein (PDB ID: 1XQ8) ^50^, an IDP, where only considered the N-terminal (residues 1–60) and NAC regions (residues 61–95), the C-terminal segment (residues 96–140), which is already disorder in the crystal structure is excluded and (10) *λ* N protein (PDB ID: 1QFQ) ^51^, an IDP protein containing 35 residues, the initial model structure of this disordered Bacteriophage *λ* N protein is taken from the RNA-unbound form of the PDB structure 1QFQ ^51^.*λ* N adopts a more structured conformation upon binding to RNA. We initiate simulations from a structured model of *λ* N in which the bound RNA is removed.

### 2.2 All atom (AA) Molecular Dynamics (MD) simulations

The GROMACS ^52^ 2018.6 package ^53^ with the Amber99sb force field (ff) ^54^ is used for our AA simulation as discussed in our earlier studies. The leapfrog algorithm is used to integrate the equations of motion. The TIP3P water model is used as the solvent. Periodic boundary conditions are applied in all three dimensions. The system is electrically neutralized by adding the required number of sodium (Na+) and chloride (Cl −) ions. The potential energy is minimized using the steepest descent algorithm ^55^. Then AA MD simulation is performed at 300K temperature and 1 atmosphere pressure, maintaining an isothermal-isobaric (NPT) ensemble. We use the Berendsen thermostat ^56^ to maintain temperature and the Parrinello-Rahman barostat ^57^ to maintain constant pressure. The Lennard-Jones (LJ) and short-range electrostatic interactions are terminated at 10 Å. We use the Particle-Mesh Ewald (PME) ^58^ method to compute the long-range electrostatic interactions. LINCS ^59^ constraints are applied to all bonds involving hydrogen atoms. We use 2 fs time step for integration. The equilibration of the system is confirmed by the saturation of the root mean square deviation (RMSD) with time. We consider the equilibrated part of the trajectory for further analysis.For GB3 and ubiquitin, we used the molecular dynamics (MD) trajectories generated in our previous study ^60^, while for homeodomain we used trajectory obtained from our earlier simulations ^61^. Additionally, we simulate GB3 using the CHARMM27 force field (ff) ^62^ while keeping all other simulation conditions identical.

### 2.3 Coarse-grained (CG) Monte Carlo (MC) simulations

The model system is simulated using the Metropolis Monte Carlo (MC) ^40^ algorithm in the canonical (NVT) ensemble. The model parameters are extracted from the AA trajectories. The system is maintained at a temperature *k*_*B*_*T* = 1, where *k*_*B*_ is Boltzmann’s constant and *T* is the absolute temperature. A total of 100,000 MC steps are performed and the equilibration is monitored by observing the potential energy of the system. Post-equilibration trajectories are used to compute various observables, including solvent distribution around bead particles and secondary structure characteristics. To improve statistical reliability, multiple (5) independent simulations with identical initial configurations are conducted and the results are averaged. We use the diameter of the solvent bead, *σ*_*s*_(= 0.25 nm), as the unit of length and the thermal energy, *k*_B_*T* (= 2.5 kJ/mol), as the unit of energy in our simulations. We fix the solvent bead density at ∼ 1 gm/cm^3^.

## 3 Analysis

We calculate the following quantities over equilibrated trajectories of the AA MD and CG MC simulations.

### 3.1 Solvent distribution function

We compute the solvent distribution function, ^63^ *ρ*(*r*) to understand the arrangement of solvent beads around solvophilic and solvophobic beads of the polymer. The *ρ*(*r*) is computed using the following function:

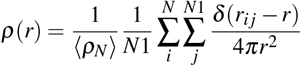

Here, ⟨*ρ*_*N*_ ⟩ represents averaged number density of solvent beads around polymer bead, i index over solvent bead, j index over reference polymer bead, *N* is the total number of solvent beads, *N*1 is the total number of solvophilic or solvophobic polymer beads, r_*i j*_ is the distance between solvent bead i and polymer bead j, 4*π*r^2^ is the surface area of a spherical shell of radius r and *δ* (*r*_*i j*_ − *r*) is the Dirac delta function.

### 3.2 Dihedral angle

We compute the phi (*ϕ*) ^64^ and psi (*ψ*) ^64^ dihedral angles per residue per frame using our in-house program.

### 3.3 Structural persistence

We calculate the structural persistence (*S*_*P*_) ^14,65^ parameter per residue using the formula:

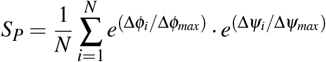

Where, *N* denotes the total number of frames. Δ*ϕ*_*i*_ and Δ*ψ*_*i*_ are the absolute values of the changes in dihedral angles *ϕ* and *ψ* of the residue in the frame *i* from the reference frame, Δ*ϕ*_*max*_ and Δ*ψ*_*max*_ are the maximum alterations possible in the Ramachandran diagram ^66^. *S*_*P*_ = 1 indicates no conformational change, whereas low *S*_*P*_ represents greater deviation from the reference structure.

### 3.4 Ramachandran plot

We generate the Ramachandran plot (RC) ^66^ using the averaged *ϕ* and *ψ* dihedral angles of residues over the equilibrated trajectory.

## 4 Results

We test our coarse-grained (CG) model on a well-structured protein, GB3 and then apply this model to other well-structured proteins, like homeodomain, ubiquitin, Chignolin, BBLand BBA and large two-domain di-ubiquitin and Adenylate kinase. We also apply our methodology to intrinsically disordered proteins (IDPs), *α*-synuclein (*α*S) and *λ* N. The resulting conformational ensembles are compared with available experimental crystal structures and all-atom (AA) simulation data.

### 4.1 Benchmark case: GB3

We perform 1 *µ*s long AA simulation for GB3 starting from the crystal structure as the initial data. The RMSD over the course of the AA simulations is shown in SI Fig. S1 (a). The data show equilibration of the system. An equilibrium snapshot from AA simulations of the GB3 protein is shown in SI Fig. S1 (b). We take the bottom-up approach, namely using the AA data to build up the CG model. We use these CG parameters to perform MC simulations on various proteins.

#### 4.1.1 Building of the CG model parameters

The COM of a residue in the crystal structure represents the center of the CG model bead. We treat the oxygen atom of the water as a solvent bead. We take polymer beads of two types: solvophilic and solvophobic, which repels the model solvent beads mimicking the hydrophilic and hydrophobic residues respectively. The beads representing the charged residues are taken to be solvophilic and the rest of the residues are taken to be solvophobic at a given pH ^67^. We use the equilibrated AA trajectory of GB3 to compute different parameters for the CG model as detailed below.

##### I. Bead size

We compute the radius of gyration, *R*_*g*_ of the protein residues based on the distances of their heavy atoms from the COM. We construct the probability distribution of the radius of gyration, *H*(*R*_*g*_) considering all the residues over the equilibrium conformations. We assign free energy corresponding to the distribution, *F*_*r*_ = −*RT* ln *H*(*Rg*), as shown in Fig. 2 (a), where, R is the ideal gas constant and T is the temperature. This profile has minima which are the stable values of the radius of gyration in equilibrium. We assign the bead radius (*σ*_*b*_/2 ∼ 0.2 nm), corresponding to the mean of the positions of the minima.

**Fig. 2.**
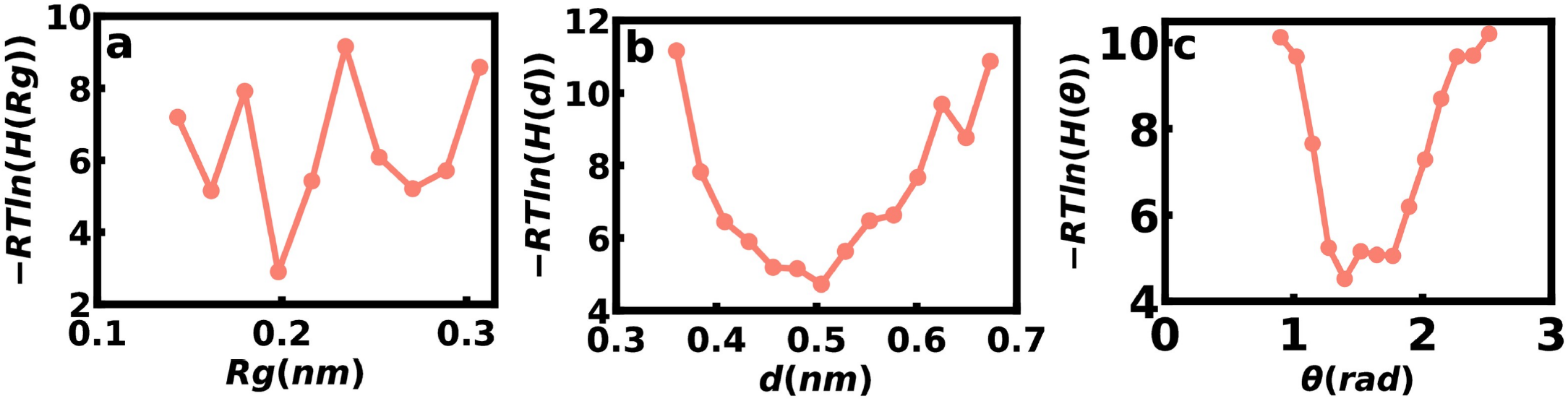
(a) Free energy profile, −*RTlnH*(*R*_*g*_) vs radius of gyration of polymer bead, *R*_*g*_, (b) Free energy profile, −*RTlnH*(*d*) vs distance, *d* between two consecutive polymer beads and (c) Free energy profile, −*RTlnH*(*θ*) vs angle, *θ* between three consecutive polymer beads.

##### II. Bonded interactions

###### Stretching potential

We compute the distribution *H*(*d*) of the bond distance, *d* is the distance between center of mass of two consecutive amino acid residues. The stretching free energy, *F*_*s*_ = −*RT* ln *H*(*d*), shown in Fig. 2 (b). The harmonic bond stretching potential between two polymer beads is modeled as 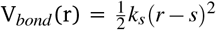, where *r* is the bond distance between two beads and *s* is the equilibrium bond distance given by the minimum of *F*_*s*_, shown in Fig. 2 (b). The stretching force constant *k*_*s*_ is calculated from the second-order derivative around the minimum of the stretching energy. Here, we find *s* = 0.5 nm and *k*_*s*_ = 2292 kJ/mol/nm^2^.

###### Bending potential

Fig. 2 (c) shows the bending free energy, *F*_*b*_ = −*RT* ln *H*(*θ*), where *H*(*θ*) represents the distribution of the bond-angle *θ* between COM of three consecutive residues in AA data. The bending potential between three consecutive polymer beads is represented by a harmonic potential: 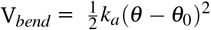, The equilibrium bond angle, *θ*_0_, is set to 1.4 rad, where *F*_*b*_ has a minimum. The bending force constant, *k*_*a*_ (87 kJ/mol/rad^2^), is calculated from the second-order derivative around the minimum of −*RT* ln *H*(*θ*) vs *θ* in Fig. 2 (c).

##### III. Dihedral interactions

The backbone dihedral angles *ϕ* and *ψ* are calculated from the atomic coordinates of the residues. We compute the joint probability distribution of the dihedral angles *P*(Γ_*i*_, Γ _*j*_). Here, Γ_*i*_ (= *ϕ*_*i*_, *ψ*_*i*_) stands for backbone dihedral angles of the *i*^*th*^ residue and Γ _*j*_ (=*ϕ*_*j*_, *ψ*_*j*_) are those for the *j*^*th*^ residue. We define the dihedral interaction profile, *F*_dih_(Γ_*i*_, Γ _*j*_) = −*k*_*B*_*Tε*_*i j*_*lnP*(Γ_*i*_, Γ _*j*_). Here *ε*_*i j*_ = 1 for *i* = *j* (intra-residual case) and the COM distance between *I* and *j*th residue (*i*. ≠ *j*, inter-residual case) is within a cut-off; otherwise, *ε*_*i j*_ = 0.

Let us first consider intra-residual distributions (*i* = *j*). We group the data into two distinct free energy profiles: one, considering the backbone dihedral angles of all the hydrophilic amino acid residues and the other, considering all the hydrophobic residues, shown in a two-dimensional grid, shown in Figs. 3 (a) and (b), respectively. This classification takes care of the chemical properties of the side chains at the simplest level, although the side chains are not explicitly considered in our model.

**Fig. 3.**
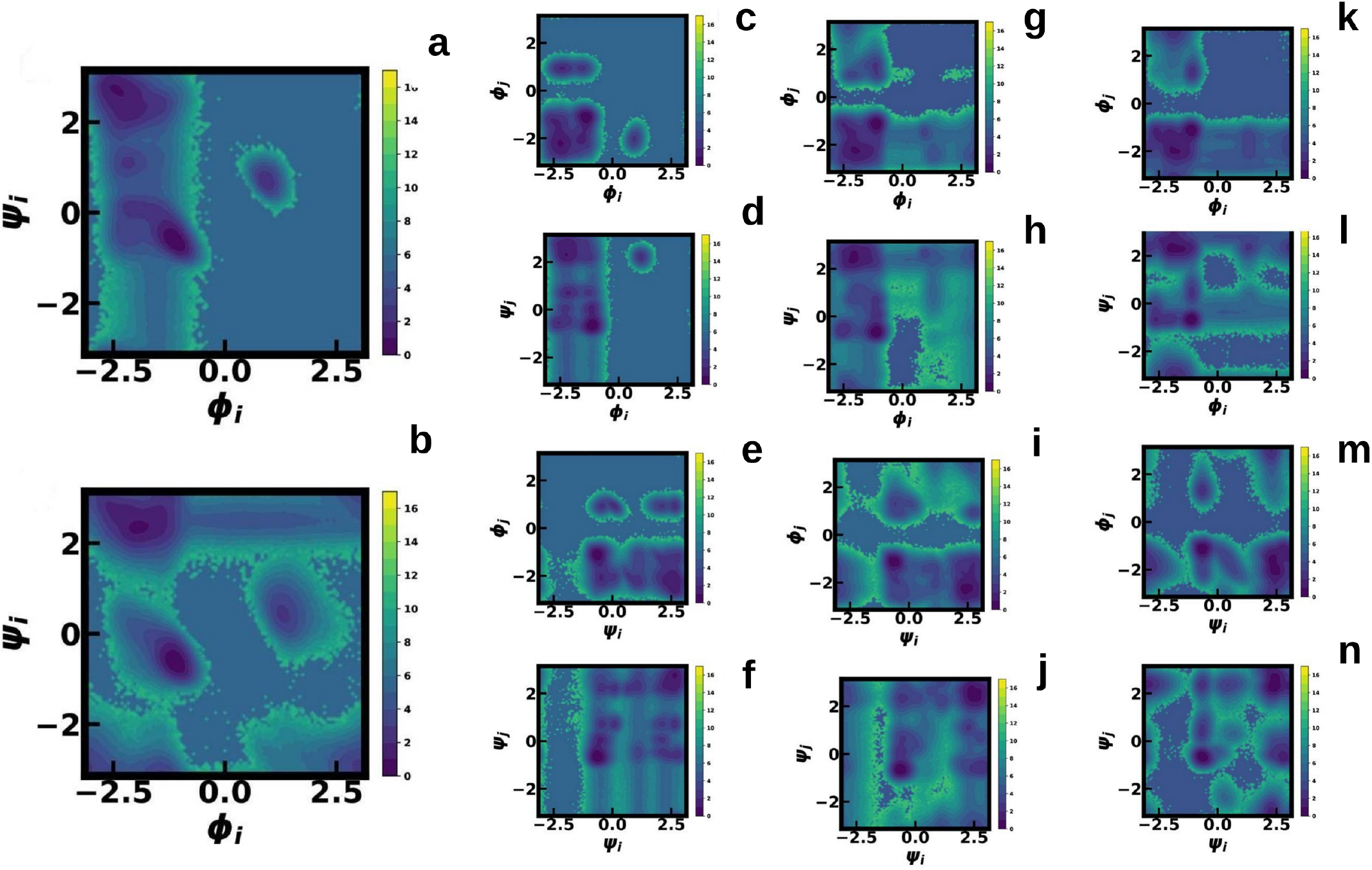
Free energy landscape for intra residual dihedral coupling, considering all (a) hydrophilic residues and (b) hydrophobic residues. Free energy landscape for inter residual dihedral coupling considering (c-f) hydrophilic-hydrophilic residues, (g-j) hydrophilic-hydrophobic/hydrophobic-hydrophilic residues and (k-n) hydrophobic-hydrophobic residues.

A similar method is adopted to calculate the free energy profile for inter-residual (*i* ≠ *j*) dihedral coupling. The dihedral angles of the corresponding residues are subsequently classified according to the hydrophilic-hydrophilic, hydrophobic-hydrophobic and hydrophobic-hydrophilic residue pairs. For each case, all possible dihedral angle combinations are considered and the negative logarithm of each joint probability distribution yields the corresponding free energy profile. The free energy landscape (FEL) plots for various combinations of dihedral angles of two different residues are shown in Figs. 3 (c)-(n).

We, thus, have the dihedral angle interaction energy landscapes classified into solvophobic and solvophilic beads corresponding to hydrophobic and hydrophilic residues respectively. Moreover, we consider (*ε*_*i j*_ = 1) the inter-residue dihedral interaction energy only when the COM of the two beads are within a cut-off 7Å. Otherwise, we ignore the dihedral interaction energy (*ε*_*i j*_ = 0). The distinction between the free energy profiles between the intra and inter-residue dihedral angles within a cut-off distance implicitly take care of the interaction between the

##### IV. Bead-solvent interaction

We compute the distribution of water oxygen atoms *ρ*(*r*), where r is the distance of oxygen and the COM of the residue. *ρ*(*r*) data are averaged over conformations considering all the hydrophilic and hydrophobic residues separately. The data - *RT* ln(*ρ*(*r*)) versus r for these two cases are shown in Figs. 4 (a) and (b), respectively. Fig. 4 (a) shows the presence of a minimum at shorter distance, while no such minimum is observed in Fig. 4(b).

**Fig. 4.**
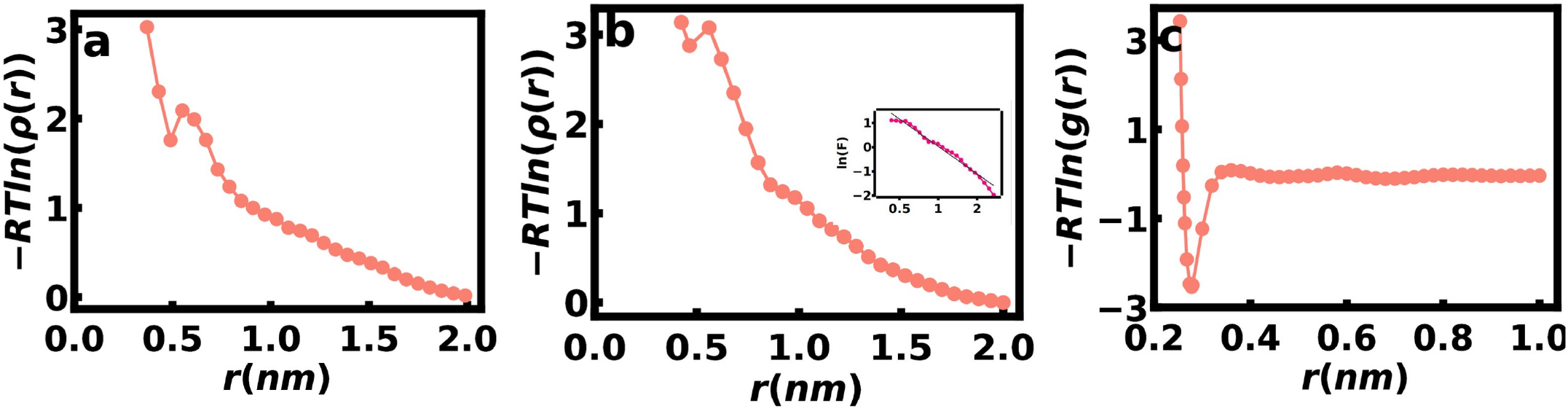
(a) Solvent density derived free energy profile, -RTln *ρ*(*r*) vs distance *r*, for water oxygen around hydrophilic residues, (b) Solvent density derived free energy profile, −*RTlnρ*(*r*) vs distance *r*, for water oxygen around hydrophobic residues, inset: Logarithmic free energy, ln *F* = −*RT* ln *ρ*(*r*), as a function of *r* for oxygen in water around hydrophobic residues and (c) Free energy profile, −*RTlng*(*r*), derived from the oxygen–oxygen radial distribution function, *g*(*r*) of water as a function of intermolecular distance, r. All the quantities are calculated from AA MD simulations data of GB3 protein.

The minimum corresponding to hydrophilic residues is described via the Lennard-Jones (LJ) 12-6 potential, defining the solvophilic polymer bead-solvent bead interaction: 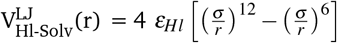 Here, *r* is the distance between the polymer bead and the solvent bead. The interaction parameter *ε*_*Hl*_ is equal to 1.25 kJ/mol, which is the minimum of the energy, *F*_*Hl*_ = −*RT* ln *ρ*(*r*) (Fig. 4 (a)). The hydrophobic beads interact with the solvent via the potential: V_Hb-Solv_(r) = 13.0 exp(−2.4*r*) (inset of Fig. 4 (b)), obtained by fitting the semilog plot of the free energy, *F*_*Hb*_ = −*RT* ln *ρ*(*r*) vs. *r*.

##### V. Solvent-solvent bead interactions

The oxygen-oxygen radial distribution function, g(r) ^63^ is calculated and the *F*_*Solv*−*Solv*_ = -*RT* ln(*g*(*r*)) versus r plot is shown in Fig. 4(c). Here, R is the distance between the center of two oxygen atoms. For solvent-solvent beads, only the non-bonded Lennard-Jones (LJ) 12-6 potential is considered: 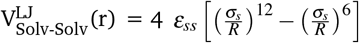 The interaction parameter *ε*_*ss*_, equal to 2.5 kJ/mol, corresponds to the minimum value of the energy, *F*_*Solv*−*Solv*_ (Fig. 4(c)).

##### VI. Non-bonded bead-bead interaction

i. Repulsive interaction: 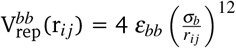, where *ε*_*bb*_ is the interaction parameter, *σ*_*b*_ is the diameter of the polymer bead and *r*_*i j*_ is the distance between two polymer beads *i* and *j*. This is to ensure that the beads cannot overlap. We set the *ε*_*bb*_ value to 2.5 kJ/mol, given by the minimum of the *F*_*r*_ (Fig. 2 (a)).
ii. Screened Coulomb interactions: 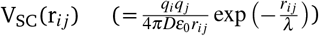, where *q*_*i*_ and *q* _*j*_ are the net charges of the residues *i* and *j*, respectively, as calculated using the pdb2pqr serve ^68^ at a given pH value. We take pH = 7 in our calculation.*ε*_0_ is the permittivity of vacuum and *D* is the dielectric constant of water (set equal to 80). Thus, medium for the electrostatic interaction is described as a continuum. We set the Debye screening length, 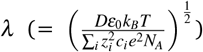 equal to 0.03 nm computed from the salt concentration and charge of the residue. Here, *e* is the elementary charge, *z*_*i*_ (= 1) the valency of *i* type of ion present in the solution, *c*_*i*_ the ionic concentration (0.1 mol L^−1^), and *N*_A_ the Avogadro’s number.

#### 4.1.2 Monte Carlo for the CG model

The CG model potential that considers different interaction terms in the system takes the following form:

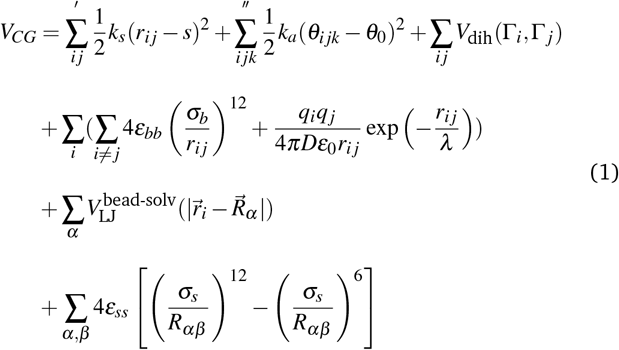

Here, *r*_*i j*_ is the distance between two polymer beads having coordinates 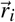 and 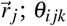 denotes the bond angle between three consecutive polymer beads *i, j*, and *k* 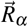 is the coordinate of the *α*^th^ solvent bead. The prime over the bonding term indicates that only two consecutive residues COM and the double prime over the bending energy term indicates that three consecutive residues are to be taken.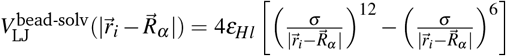 for solvophilic beads and 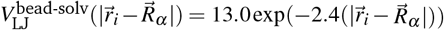 for solvophobic bead at 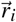 and solvent bead 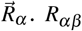 is the distance between two solvent beads are having coordinates 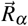 and 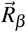 The bead-dihedral interactions are implicitly accounted for by bead-bead distance cut-off where the dihedral interactions are considered for beads within the cut-off distances. For the inter-bead dihedral contribution, we consider only those beads that are within 7.0 of each bead in every frame. The bead-solvent and solvent-solvent interactions are truncated at a cutoff distance of 6.25.

We take all the parameters in the model potential as determined from the GB3 AA trajectory, given in Table 1. The initial structure is taken from the crystal structure of GB3. Each bead is located at the center of mass of a residue determined from the heavy atoms in a given residue. The beads are classified as either solvophilic or solvophobic, as given in SI Table S1, depending on hydrophilic or hydrophobic residue obtained from water oxygen distributions. The initial values of the dihedral angles of different residues are taken from the crystal structure and the solvent molecules are the oxygen atoms of water molecules in an equilibrated AA configuration . The coarse-grained (CG) polymer and solvent beads are placed in a cubic simulation box of length L=5.6 nm, with the periodic boundary conditions applied in all directions (x, y and z). We use 56 polymer beads and 5,444 solvent beads in our simulation.

**Table 1.** Table of CG parameters.

| Parameter | Value |
| --- | --- |
| $\sigma_s$ | 0.25 nm |
| $k_s$ | 2292 kJ/mol/nm<sup>2</sup> |
| $s$ | 0.5 nm |
| $k_a$ | 87 kJ/mol/rad<sup>2</sup> |
| $\theta_0$ | 1.5 rad |
| $\epsilon_{bb}$ | 2.5 kJ/mol |
| $\sigma_b$ | 0.4 nm |
| $\epsilon_{HI}$ | 1.25 kJ/mol |
| $\epsilon_{ss}$ | 2.5 kJ/mol |

During a MC move, a bead, either polymer or solvent, is selected at random. All three position coordinates along with two dihedral angles are given a random movement to generate a new configuration of the selected polymer bead. For a solvent bead, only three position coordinates are updated. The energy changes due to the change in the degrees of freedom of the bead are computed based on various interactions in the CG potential, *V*_*CG*_. The contribution due to changes in the dihedral angles, *V*_dih_(Γ_*i*_, Γ _*j*_) is computed as follows. The intra-bead (i=j) interaction between the dihedral of the beads is calculated by linear interpolation from the two-dimensional free energy profile, F grids in Fig. 3. We choose the type of grid depending on if the bead is solvophilic or solvophobic. The inter-bead (i≠ j) dihedral coupling depends on the solvophilic or solvophobic character of the pair of beads. We use the *V*_dih_(Γ_*i*_, Γ _*j*_) parameters shown in Fig. 3 to interpolate.

We allow the system to equilibrate, which is monitored via the total energy (SI Fig. S2) and calculate the structural quantities over the equilibrated trajectory. We calculate the solvent bead distributions *ρ*(*r*) at a distance *r* from the bead center around solvophilic and solvophobic beads (Fig. 5 (a)) in the scaled unit. Solvophilic beads form strong interactions with solvent beads, leading to the formation of a well-defined first solvation shell. As a result, the local solvent density increases near the reference bead, producing a pronounced first peak in solvent distribution. In contrast, solvophobic beads do not interact strongly with the solvent and therefore the solvent distribution near the bead is not significantly enhanced; consequently, the first peak is weak or absent. In the inset, the corresponding AA data are shown. For the AA results, the x-axis represents the distance between the COM of the given residue (hydrophilic or hydrophobic) and oxygen atom of water particles in nm, while the y-axis represents the normalized average solvent number density at distance *r*, which is also dimensionless. Similar to the CG solvophilic beads, hydrophilic residues in the AA data form strong interactions with solvent molecules and stabilize structured hydration shells, leading to a clear first peak in the solvent distribution. On the other hand, hydrophobic residues interact weakly with solvent molecules, so the first peak in the solvent density is weak or absent. Overall, the CG results reproduce the qualitative trend observed in the AA data. We further compute structural persistence, S_*P*_ per residue in equilibrated CG MC and AA MD trajectories, as shown in Fig. 5 (b). The CG and AA simulation data for S_*P*_ exhibit good agreement.

**Fig. 5.**
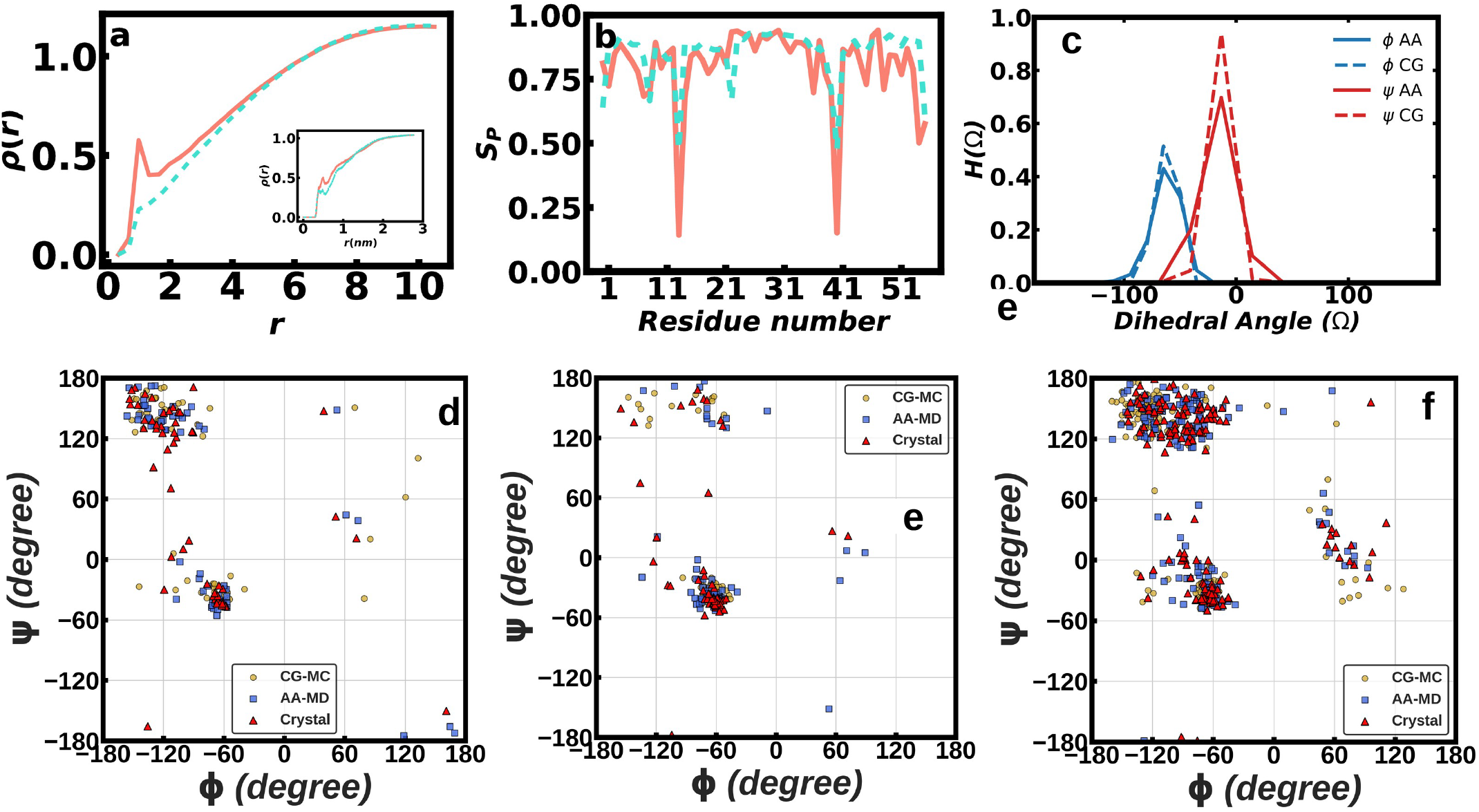
(a) Solvent distribution, *ρ*(*r*), around solvophilic (salmon, solid line) and solvophobic (turquoise, dashed line) beads in the equilibrated CG MC trajectory of GB3. The inset shows the solvent distribution around hydrophilic (salmon, solid line) and hydrophobic (turquoise, dashed line) residues in the equilibrated AA MD trajectory. (b) Structural persistence (S_*P*_) per residue in equilibrated CG simulations (salmon, solid line) and AA MD simulations (turquoise, dashed line). (c) Probability distributions of the *ϕ* and*ψ* dihedral angles of LYS10 in the GB3 protein, calculated from equilibrated trajectories. Solid linesrepresent AA MD data, while dashed lines represent CG MC data. Blue denotes the *ϕ* dihedral angle, whereas red denotes the *ψ* dihedral angle. Comparison of Ramachandran (RC) plots for (d) GB3, (e) homeodomain, and (f) di-ubiquitin proteins obtained from the crystal structure, the final structure from all-atom molecular dynamics (AA MD) simulations, and the final structure from coarse-grained Monte Carlo (CG MC) simulations.

**Fig. 6.**
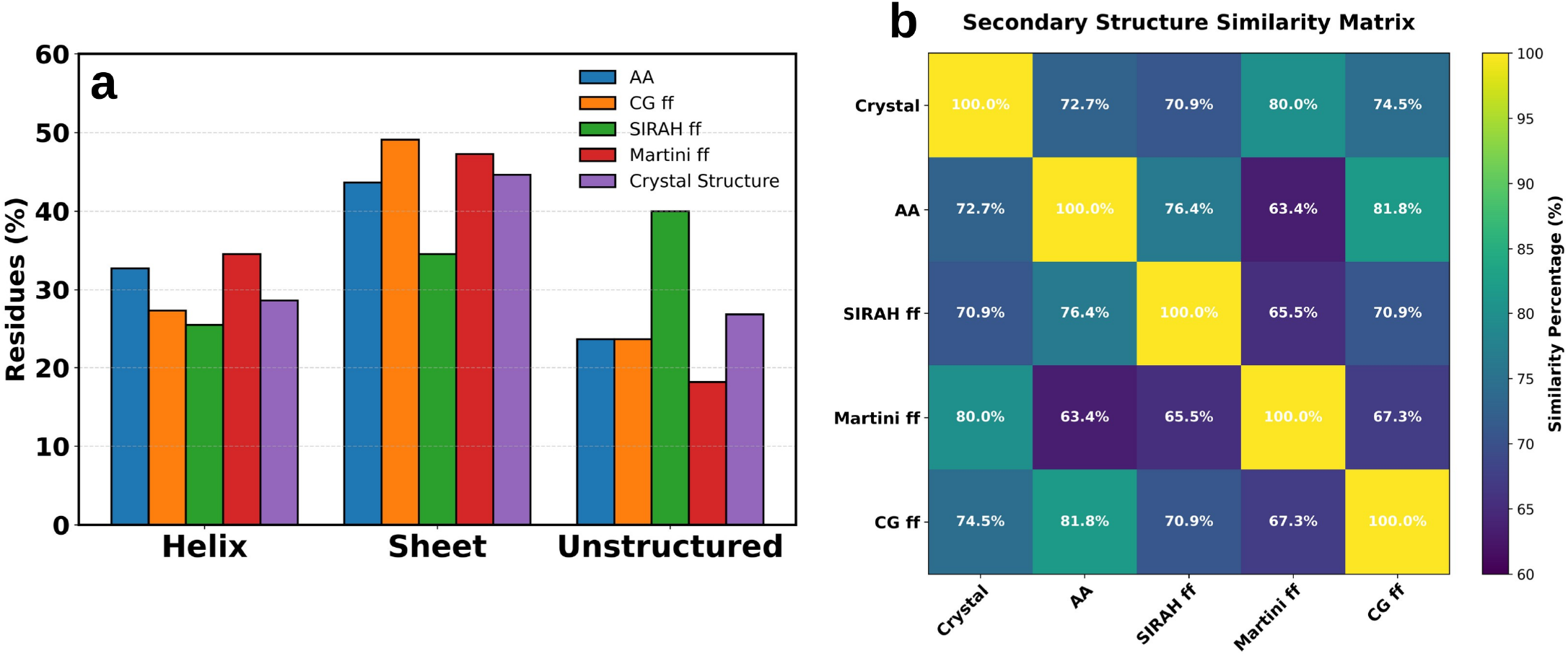
(a) Bar plot showing residue counts in % in helix, sheet, and unstructured regions for five cases—AA simulation (AA), our structure-based CG force field (CG ff), SIRAH, Martini, and the crystal structure. (b) Secondary-structure similarity matrix, where off-diagonal elements quantify agreement between methods.

In Fig. 5(c), we show the probability distributions of the *ϕ* and *ψ* dihedral angles for randomly chosen LYS10 in equilibrated trajectories. The solid line represnt AA distribution while dashed denote CG distribution. The distributions from AA MD and CG MC simulations show good agreement. Dihedral distributions for a few additional residues are shown in SI. Fig.S3. We examine how far the structural aspects are captured in our CG simulations. To this end, we compare the secondary structure preferences of each residue from the CG MC simulations. We assign the secondary structure elements to each bead, namely, helix (H), *β* -sheet (S) and unstructured (U) depending on the values *ϕ* and *ψ* for each conformation ^69^. The unstructured (U) category includes all conformations that do not fall into the helix (H) or *β* -sheet (S). We assign a given bead the preferred structural element, the one that occurs the maximum number of times over the equilibrium trajectory. A similar strategy is followed for the AA data and the secondary structural elements are assigned directly from the dihedral angles of the backbone. SI Table S2 presents the structural preferences of each residue in CG MC simulation, along with their corresponding secondary structures in the crystal structure and AA MD simulations. We find that the secondary structural elements show ∼80% agreement with the residues observed in the crystal structure and AA data.

We also analyze the Ramachandran (RC) plots to compare the final structures obtained from the CG simulations with those from the AA simulations and the crystal structure of the GB3 protein. The RC plots, shown in Fig. 5(d), confirm excellent agreement among the three structures. We show in Table 2 comparison of percentages of residues in a given structural element over the CG and AA trajectories along with those in the crystal structure. the helix (H) elements are in excellent agreement to the AA and crystal structure data. Our model slightly over-estimates the *β* -sheet (S) compared to the crystal structure, while the AA data underestimates the *β*-sheet (S) structure. The unstructured (U) region in the CG model agrees better with the crystal structure compared to the AA data. Overall, our model reproduces the secondary structural elements as accurately as in the AA data.

**Table 2.**
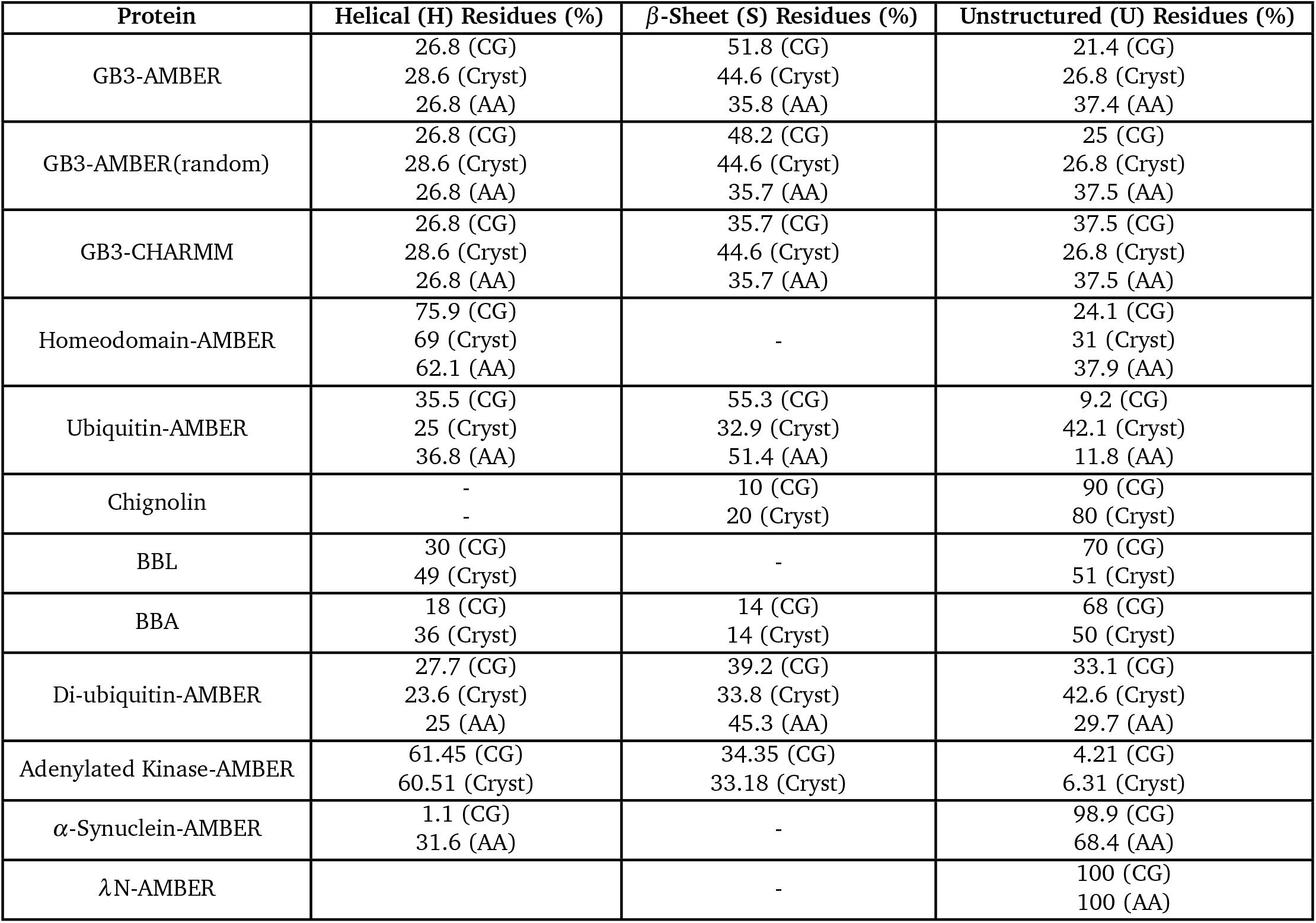
Comparison of secondary structure preferences of proteins. AA denotes trajectory of all atom molecular dynamics simulations and CG denotes conformations based on coarse-grained Monte Carlo simulations. ‘H’ corresponds Helix, ‘S’ corresponds sheet and ‘U’ corresponds to other than element helix or sheet i.e. loop/coil/turn/bend region of the protein.

We carry out controlled studies to check the robustness of our model. It is unclear whether the good agreement between the CG and crystal structures arises from any bias introduced by our initial choices. To investigate this, for GB3, we add to the crystal dihedral angles random number within a range of −*π*/10 to +*π*/10. We perform the CG simulations starting from the random structure. Here also, the resulting residue-wise secondary structure matches well with the crystal structure (80.3%) (SI Table S3). As shown in Table 2, our CG model reproduces the percentages of different structural elements in very good agreement to the AA and the crystal structure data.

We examine whether using the AA trajectory introduces any force field bias in the CG parameters. To check this, we perform CG simulations on GB3 model system using interaction parameters derived from widely used AA force field, CHARMM and compare the resulting structural features with crystal data and those obtained from corresponding AA MD simulations. The data are shown in SI Tables S4. The structural agreement observed in all cases is consistent with our previous findings based on the AMBER force field, showing >=75% similarity with the crystal and AA data (SI Tables S4), along with similar percentages of different secondary structural elements (Table 2).

### 4.2 Transferability of the GB3 CG model parameters to other well structured proteins

Next, we check how far the structural data for other well-structured proteins can be reproduced by using the CG parameters derived from GB3 AA trajectory. We consider several other proteins having well defined experimental structure: homeodomain, Ub, chignolin, BBA and BBL. We verify that the CG interaction parameters derived from AA trajectories for homeodomain and Ub proteins are similar to those of GB3. This similarity can be traced to the fact that the CG parameters are taken as an average over the entire protein and the beads are distinguished based on gross feature, namely, solvophobic and solvophilic interactions. We carry out CG simulation using the GB3 CG parameters for all these proteins and compare the structural data for homeodomain and Ub from CG simulations, Crystal structures and AA simulations Only CG simualtions for the remaining proteins are performed and compared to the crystal structures. The charged residues in these proteins are taken to be solvophilic, while the other beads are classified to be solvophobic.

#### Homeodomain

At first we consider protein homeodomain which have well defined crystal structure. It is a small, 58-residue protein that binds to specific DNA sequences and functions as a transcription factor, playing a crucial role in gene regulation. SI Table S5 shows the secondary structural elements of each residue derived from the crystal structure, AA simulations and CG MC simulations for homeodomain. We observe good agreement: The CG MC simulation reproduces the crystal structural data with 90% accuracy (SI Table S5). The residue-wise secondary structural elements from CG MC simulations agree well with the AA MD results, showing an agreement of 81% (SI Table S5).The RC plot based on final structure is depicted in Fig.5(e). Our CG model captures the secondary structural elements as good as the AA data and are quite comparable to the crystal structure data (Table 2).

#### Ubiquitin

Ubiquitin is a 76 residue globular protein involved in the targeted degradation of cyclins and other regulatory proteins. In case of ubiquitin, we find that both the AA and CG data overestimates the structured portion of the protein, while underestimating the unstructured parts. SI Table S6 shows detailed comparison: The secondary structural elements show 67% matching with crsytal structure. Although this agreement is poorer compared to other proteins, the mismatched residues mostly correspond to the unstructured (U) conformation in the crystal structure. The residues adopting structured conformations (H or S) determined from CG simulations agree well with the crystal structure. Compared to the AA MD simulation data (also in SI Table S6), the agreement improves significantly to 92%. Table 2 lists the agreements secondary structural element wise: Here also, the structure elements in particular H conformation is overestimated in both AA and CG models, while the U elements are underestimated compared to the crystal structure. We further show the RC plots based on the final structures obtained from the CG model and compare them with the AA and crystal structure data (see SI Fig. S4). The good agreement in the RC plots indicates the accuracy of the model in reproducing the backbone conformational preferences and, in particular, in preserving the secondary structure propensities observed in the atomistic and experimental structures.

#### Chignolin

We study another small protein, Chignolin ^70^, which contains ten amino acid residues. Chignolin is widely used in computational biology and biochemistry as a benchmark model for studying protein-folding mechanisms. Here, we compare residue wise secondary structure data with crystal structure and summarize the results in the SI Table S7. The CG data show good agreement with the crystal structure, with an accuracy of 90%. In Table 2, we summarize the percentage of residues adopting different secondary structure, H or S in both the crystal structure and CG simulation.

#### BBL protein

We next apply our method to the BBL protein ^71^, a small folded protein containing forty seven amino acid residues. BBL is frequently used in structural biology as a model system to understand how proteins fold into their three dimensional structures. The residue-wise secondary structure analysis, summarized in SI Table S8, shows that the CG data exhibit 77% agreement with the crystal structure. We also summarize in Table 2, the percentage of residues adopting different secondary structure elements in both the CG simulations and the crystal structure.

#### BBA protein

Next, we apply our CG model to study another small folded protein, BBA ^72^, which contains twenty eight amino acid residues. BBA is also frequently used in computational biology and biochemistry as a model system for studying protein folding mechanisms. SI Table S9 shows the residue wise secondary structure assignments for both the CG simulations and the crystal structure. The CG data show good agreement with the crystal structure with an agreement of 75%. The percentage of residues adopting different secondary structure elements in the crystal structure and CG simulations is summarized in Table 2. As Chignolin, BBL and BBA are well studied folded proteins, we validate our results by comparing them with the corresponding crystal structures only.

### 4.3 Cases of multi-domain proteins

Next we extend our CG model with the GB3 parameters to study conformational preference for two-domain proteins: (i) Di-ubiquitin a 148-residue two-domain protein that serves as a signal for proteasomal degradation (see Fig.1(e)). Two ubiquitin domains are connected by a covalent isopeptide bond where the C-terminal glycine (Gly76) of the first ubiquitin is linked to lysine (Lys48) of the second ubiquitin. and (ii) adenylate kinase (see Fig.1(k)), comprising of two domains and 428 amino acid residues, that maintains cellular energy homeostasis by catalyzing the conversion of two ADP molecules into ATP and AMP and undergo large-scale conformational transitions during catalysis. In case of di-ubiquitin, the CG parameters are computed from AA trajectory and agree well to those for GB3. We use the GB3 CG parameters to both the proteins.

We compare residue-wise secondary structure preferences obtained from the CG simulations with those derived from the crystal structure and AA simulations for di-ubiquitin (SI Tables S10 (a) & (b)). The CG results show good agreement with the crystal structure. Approximately 70% of residues exhibit matching secondary structure preferences. Most of the mismatched residues are having a U conformation in the crystal structure. Compared to the AA MD data, approximately 74% of residues show consistent secondary structure preferences, while the remaining residues predominantly adopt S or U conformations (Table 2). The corresponding RC plot based on final structure is depicted in Fig.5(f). Table 2 shows a comparative analysis of percentage of secondary structural elements of di-ubiquitin. The data show that the agreement of the secondary structural elements is quite good for all the cases. In case of adenylate kinase, the RC plot based on the final structure is shown in SI Fig. S4. The results indicate good structural agreement between the CG model and the crystal structure. Table 2 further shows comparative analysis of secondary structure element. Overall, these results demonstrate that the present CG model robustly preserves secondary structure features and captures the global structural integrity of multidomain proteins as well.

### 4.4 Case of intrinsically disordered protein (IDP)

Now we consider the cases of couple of IDPs: *α*S and *λ* N. *α*S is a small abundant neuronal protein in the brain, best known for its role in synaptic function and pathological aggregation in Parkinson’s disease. *λ* N is an anti-termination protein that recognizes the nut site on the phage RNA and enables transcription to continue past termination signals. We compute all bonded and non-bonded interaction parameters from the AA MD trajectory of *α*S, the initial structure of *α*S being taken from the crystal structure deposited under PDB ID: 1XQ8. We find that the bond force constant (*k*_*b*_ = 746 kJmol^−1^ nm^−2^, as detailed in the SI) and the angle force constant (*k*_*a*_ = 30 kJmol^−1^ rad^−2^, as detailed in the SI) are smaller than those obtained for GB3, unlike the cases of well-structured proteins. The remaining parameters are comparable. We observe similar elastic constants from AA trajectories of *λ* N. These reduced elastic constants reflect the increased conformational flexibility inherent to the intrinsically disordered nature of the *α*S protein. This also illustrates the importance of elastic properties of the backbone for secondary structures. We perform CG simulations of *α*S and *λ* N using the force constants calculated for *α*S. Since the IDPs lack well defined crystal structure, we compare the structural elements from our CG simulations to those from AA data only.

*αS:* We show in SI Tables S11 (a) & (b) comparison between CG and AA MD data for *α*S. Approximately 67.4% of residues exhibit consistent secondary structure preferences (SI Tables S11 (a) & and (b)). In this case, the unmatched residues in the CG simulations predominantly sample U conformations, whereas they adopt H conformations in the AA simulations. Table 2 shows the overall percentages of the secondary structure elements in the CG and AA data. The U elements are in vast majority in CG structures than the AA data. The lack of structure is consistent with the intrinsically disordered nature of the protein which is better reproduced in the CG simulations than that in the AA simulations.

*λ N*: In case of *λ* N, we observe that in both CG and AA simulations, the residues preferentially adopt U conformations (Table 2 and SI Table S12) similar to *α*S.

## 5 Discussions

### 5.1 Comparison to Martini and SIRAH Models

An explicit comparison of our model to other widely used CG models is worthwhile here. We perform to this end simulations of the GB3 protein using both the SIRAH and Martini CG force fields in explicit solvent. The details of these simulation methods are provided in the SI. The final GB3 structures after 1 µs simulations using SIRAH and Martini CG force fields are shown in SI Fig.S5 (a) and (b), respectively, with residues colored by secondary structure (helix, sheet, unstructured). Next, we count the secondary-structural elements based on final structure for four cases: AA simulation, our CG force field, SIRAH, Martini, as well as experimental crystal structure—to provide a clear basis for comparison (see Fig.6a). The CG force field SIRAH exhibits a bias toward disorder, underestimating helices and sheets. In contrast, the Martini force field shows the opposite tendency i.e. over-stabilizing ordered secondary structure and significantly under predicting unstructured structures compared to experiment. on the other hand, our CG model is able to show results closer to both AA and crystal structure data.

We further construct similarity matrix by performing pair wise residue-by-residue comparison of secondary structure assignments (helix, sheet or unstructured) between all five systems (Fig.6(b)). For each pair of system, the percentage similarity has been calculated as the number of residues with identical secondary structure assignments divided by the total number of residues, multiplied by 100. It gives a symmetric matrix with 100% self-similarity on the diagonal and off-diagonal elements quantify the agreement between different methodologies. Our CG model shows strongest overall performance, achieving the highest pairwise similarity with the AA simulation (81.8%), indicating it most faithfully reproduce the structural details captured by the AA reference. Our CG model also maintain a good agreement with experimental crystal structure as well (74.5%). On the other hand, Martini shows relatively good agreement with the crystal structure (80.0%) but poor agreement with AA (63.4%) and SIRAH (65.5%). Conversely, SIRAH’s strongest agreement is with AA (76.4%), but it shows lower similarity to crystal (70.9%) and the lowest agreement with Martini (65.5%). The matrix thus quantitatively confirm that our CG force field simultaneously preserves AA level structural detail and maintains good agreement with the experimental fold. It therefore provides the most balanced and reliable residue-level secondary structure representation among the tested CG models.

### 5.2 Advantages of the proposed CG model

Earlier CG models for DNA–protein systems ^73,74^ and DPG-MC approach to generate polypeptides ^36^ and protein structures ^37^ heavily rely on data bank structures. Such approaches have inherent limitations: The data bank do not generate information on interaction between water and the bio-molecules, although these interactions play primary roles in adopting bio-molecular structure. The coarse-graining requires statistical averages over a wide range of conformations to accurately derive effective interaction potentials. It is difficult to obtain adequate averaging using limited number of available data bank structures. Moreover, the PDB structures lack consistent ensemble information and may be influenced by crystallization artifacts. Our current CG model surpasses these limitations by deriving the CG parameters from AA trajectories. This ensures that the solvent interactions are taken care of, the thermodynamic ensemble is well defined and there are sufficient conformation to extract meaningful CG parameters. We find, moreover, that the CG parameters obtained from different AA models can reproduce the structure with comparable accuracy.

It is important to point out that our model is capable to reproduce the structural elements in good agreement to the crystal structure and AA data where the input parameters are not sensitive to the secondary structural elements of the residues. The backbone elastic energy, dihedral and bead-solvent interactions play key role to stabilize the secondary structural elements. Our computational model offers several advantages compared to other CG models reported in the literature: (i) It is a single-site meso-scopic model so that it can be easily applied to large proteins and multiple proteins in a solvent. (ii) In each MC step, both Cartesian coordinates and backbone dihedral angles are updated simultaneously, avoiding separate Cartesian and dihedral-space sampling schemes and resulting in improved computational efficiency. (iii) It employs an explicit-solvent representation and distinguishes the beads based on solvent interaction. (iv) The backbone hydrogen bonding interactions are incorporated implicitly through the CG interaction profiles derived from atomistic simulation trajectories.

The CG force field parameters derived from GB3 are transferable to other structured proteins. However, when we use the stretching and bending force constants derived from the GB3 protein, we observe lack of equilibration in both IDP proteins. This suggests that the backbone flexibilities play key role in adopting the equilibrium structure. On the other hand, the CG parameters for one IDP work well for another IDP. The transferability of the system arises from two main reasons: (1) The chemical environment of the backbone formed from peptide bonds is similar for different proteins with comparable elastic constants. (2) A broad categorization of solvent and dihedral interactions based on hydrophilic and hydrophobic residues is sufficient to describe the protein secondary structural elements. However, it must be pointed out that we have tested transferability only for restricted number of cases. A general proof of transferability is quite difficult. The main importance of our study is as follows: Given AA trajectory for a protein, we show a method to carry out systematic coarse graining while retaining the structural information. Based on CG model the phase behaviour and dynamics of an aqueous suspension of proteins can be studied which is beyond the scope of AA simulations.

### 5.3 Limitations and Future Directions

Our model can be improved in several important ways: (1) The CG interaction parameters may be improved by considering a family of proteins instead of a single protein as done in our studies. We consider a single protein data to illustrate the working of the proposed CG model. (2) The electrostatic interaction effects are included as a constant with dielectric constant *ε*, considering the medium as a continuum. On the other hand, we use a monoatomic water model, where each oxygen atom of a water molecule is treated as a solvent bead to calculate the solvent-solvent and bead-solvent interactions. It is important to treat all the interactions on equal footing. In particular, taking the dipole moment of water molecules explicitly into account would lead to a more realistic solvent model instead of using dielectric continuum. (3) Inclusion of side chain information is needed to describe ligand binding interactions through side chains. (4) Another limitation of the current implementation is that the simulations are performed using an in-house code, which has limited efficiency for very large bio-molecular systems. Interfacing the model with established molecular simulation software would significantly improve computational performance and scalability. This will be an important direction for future development of the model to tackle diverse situations.

## 6 Conclusion

To summarize, we show that the secondary structural elements of the protein can be captured by introducing appropriate CG model potential energy with backbone elasticity, dihedral interaction and distinction of solvent interactions with the solvopobic and the solvophilic beads. The CG force-field parameters are derived from AA simulation trajectories. This model would enable to perform much faster simulations for studying phenomena involving multiple protein molecules, such as protein aggregation. Our CG model needs to be improved to incorporate side chain dihedrals and water dipoles moment and can be extended study the in bio-molecular systems in terms of the CG variables. Similar CG model may extended to other bio-molecular systems.

## Data availability

We have used GROMACS 2018 MD simulation package to perform all the atomistic simulations. The software can be found at https://manual.gromacs.org/documentation/respectively. The coarse grained monte carlo simulation has been using our in house code available at https://github.com/snbsoftmatter/mc-protein-cg.

## Author contributions

KK and AGM: Conceptualization, data curation, formal analysis, methodology, validation, visualization, software, investigation, writing – original draft, writing – review & editing. JC: conceptualization, methodology, validation, project administration, resources, supervision, writing – review & editing.

## Acknowledgements

The authors thank the Technical Research Centre, S. N. Bose National Centre for Basic Sciences for computational support. K.K. and A.G.M. thank Anirban Paul for helpful discussions. K.K. thanks the University Grants Commission (UGC), India [F. No. 16-6(DEC. 2018)/2019(NET/CSIR)], and A.G.M. thanks DST, India, for an INSPIRE fellowship (IF170961) as financial support. JC acknowledges financial support from the CSIR Emeritus Scientist scheme.

