## Supplementary material for "Simulation of Protein Structure using a Coarse-Grained Potential incorporating the Backbone Dihedral Interactions": SI file

### Supporting information text:

#### I. Bonded interaction parameters of $\alpha$-synuclein

(i) Stretching potential: We follow the same procedure as used for the GB3 system. Specifically, we compute the distribution $H(d)$ of the bond distance $d$, defined as the distance between the centers of mass of two consecutive amino acid residues. The corresponding stretching free energy is obtained as $F_s = -RT \ln H(d)$, as shown in SI Fig. S4(a). The harmonic bond-stretching potential between two polymer beads is modeled as

$$V_{\text{bond}}(r) = \tfrac{1}{2} k_s (r - s)^2,$$

where $r$ is the bond distance and $s$ is the equilibrium bond length, taken as the minimum position in $F_s$ (SI Fig. S4(a)). The stretching force constant $k_s$ is determined from the second derivative of $F_s$ at this minimum. Using this procedure, we obtain $s = 0.43$ nm and $k_s = 746\ \text{kJ}\,\text{mol}^{-1}\,\text{nm}^{-2}$.

(ii) Bending potential: We follow a procedure similar to that used for the GB3 system. SI Fig. S4(b) shows the bending free energy, $F_b = -RT \ln H(\theta)$, where $H(\theta)$ denotes the distribution of the bond angle $\theta$ formed by the centers of mass of three consecutive residues in the AA data. The bending potential between three consecutive polymer beads is modeled using a harmonic form,

$$V_{\text{bend}} = \tfrac{1}{2} k_a (\theta - \theta_0)^2.$$

The equilibrium bond angle $\theta_0$ is set to 1.6 rad, which corresponds to the minimum of $F_b$. The bending force constant $k_a$ ($30\ \text{kJ}\,\text{mol}^{-1}\,\text{rad}^{-2}$) is obtained from the second derivative of $-RT \ln H(\theta)$ with respect to $\theta$ near this minimum, as shown in SI Fig. S4(b).

#### II. Details of CG simulation using SIRAH forcefield:

To perform the coarse-grained (CG) simulations using the SIRAH force field,[1] the initial crystal structure of GB3 (RCSB PDB ID: 2OED)[2] was obtained from the Protein Data Bank. All simulations were performed using the GROMACS[3] package with explicit water. First, the PDB2PQR[4] server was used to assign the correct protonation states assuming neutral pH, following the AMBER naming scheme. After mapping the system into the CG representation, the protein was solvated in water box of dimension 7.38893x6.96635x6.03303 $nm^3$ using the SIRAH WT4 water model.[5] The system was then neutralized and maintained at physiological salt concentration by adding 29 $Na^+$ and 27 $Cl^-$ ions, respectively. Finally the total number of CG beads including solvent was 3849.

Next, the system was energy-minimized in a two-step manner. First, the protein side chains were energy-minimized for 50,000 steps using the steepest descent algorithm while restraining the backbone. Thereafter, the entire system was energy-minimized for 5,000 steps. In the next step, the solvent molecules were equilibrated around the protein for 5 ns while applying harmonic positional restraints on all CG beads. The system temperature was maintained at 300 K using the V-rescale thermostat. To ensure better solvation of the protein side chains, the system was further equilibrated for 25 ns. Finally, a production run of 1 $\mu$s was performed while maintaining the pressure at 1 atm using the Parrinello–Rahman barostat with isotropic pressure coupling. The integration time step for all simulations was fixed at 20 femtoseconds.

To calculate the secondary structure of each amino acid in the final structure obtained from the production run, the sirah-ss tool from the SIRAH tools[6,7] was used. This tool assigns secondary structure based on hydrogen-bond–like interactions and the instantaneous values of the backbone torsional angles.

#### III. Details of CG simulation using Martini forcefield:

The Martini[8] CG simuilation has been performed in GROMACS[3] simulation package considering initial structure 2OED[2] from RCSB PDB data bank.The coarse-grained molecular dynamics simulations were carried out using the Martini3 virtual-site implementation of an enhanced GōMartini model.[9] Construction of the Gō-like network in the GōMartini framework needs a contact map based on the initial atomistic input structure. This contact map was generated using the GoContactMap web server (http://pomalab.ippt.pan.pl/GoContactMap) with default settings. In total, 8875 CG water beads and 99 sodium ions along with 97 cl itoms (for neutralization of the charge of the simulation box and maintain 0.15M salt concentration) were added to the system. After energy minimization, the system was simulated for 1$\mu$s in the NPT ensemble at 300K and 1bar using V-rescale thermostat and Parrinello-Rahman barostat respectively. All coarse-grained simulations followed the recommended simulation parameters, including a 20 fs time step.

The secondary structure has been estimated based on final structure obtained after production run. At first, final structure was backmapped using CG2AT modules.[10] Secondary structure in the backmapped Martini structures was identified using a Ramachandran-based STRIDE-like classification of instantaneous $\phi$–$\psi$ angles, following the same criteria as implemented in the sirah-ss tool for SIRAH simulations.[6,7]

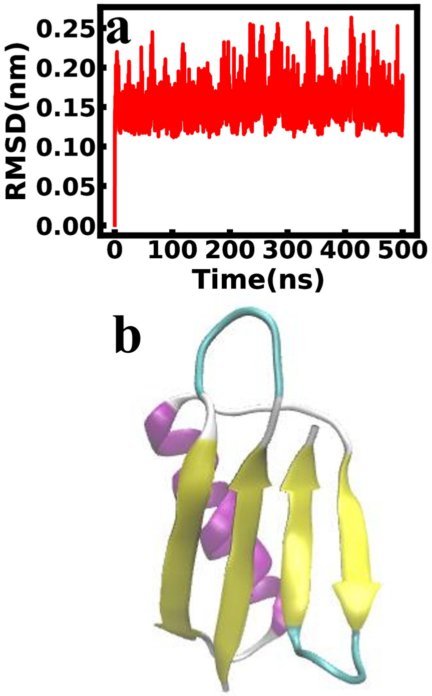


Figure S1: (a) RMSD over the course of the simulation and (b) an equilibrated snapshot of the protein GB3 at 500 ns time span in AA simulation.

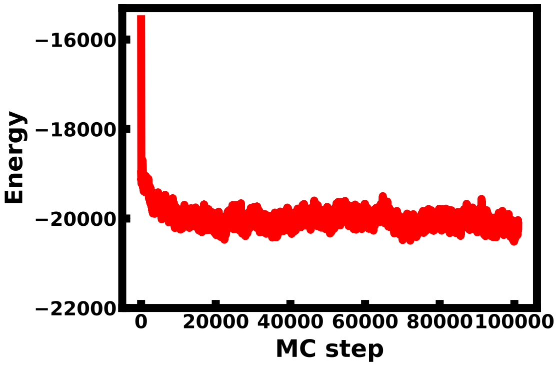


Figure S2: Total system energy at each Monte Carlo step during coarse-grained simulations of GB3.

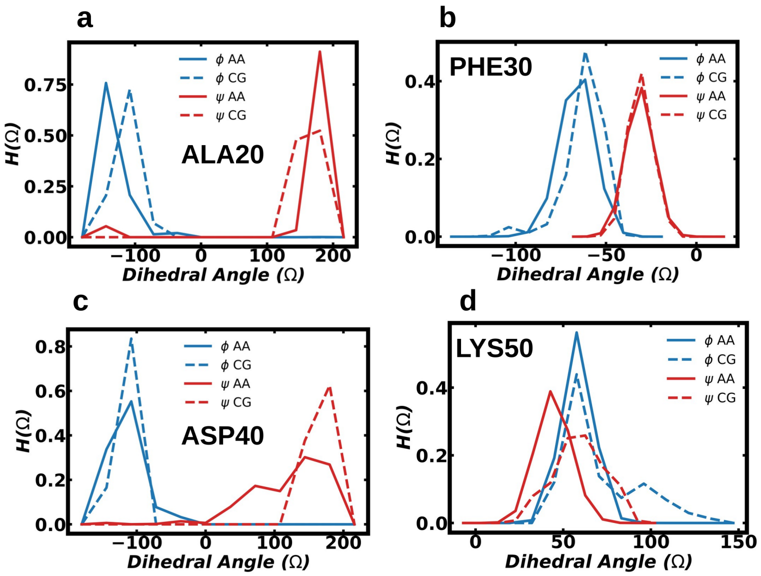


Figure S3: Probability distributions of the $\phi$ and $\psi$ dihedral angles of (a) ALA20, (b) PHE30, (c) ASP40 and (d) LYS50 in the GB3 protein, calculated from equilibrated trajectories. Solid lines represent AA MD data, while dashed lines represent CG MC data. Blue denotes the $\phi$ dihedral angle, whereas red denotes the $\psi$ dihedral angle.

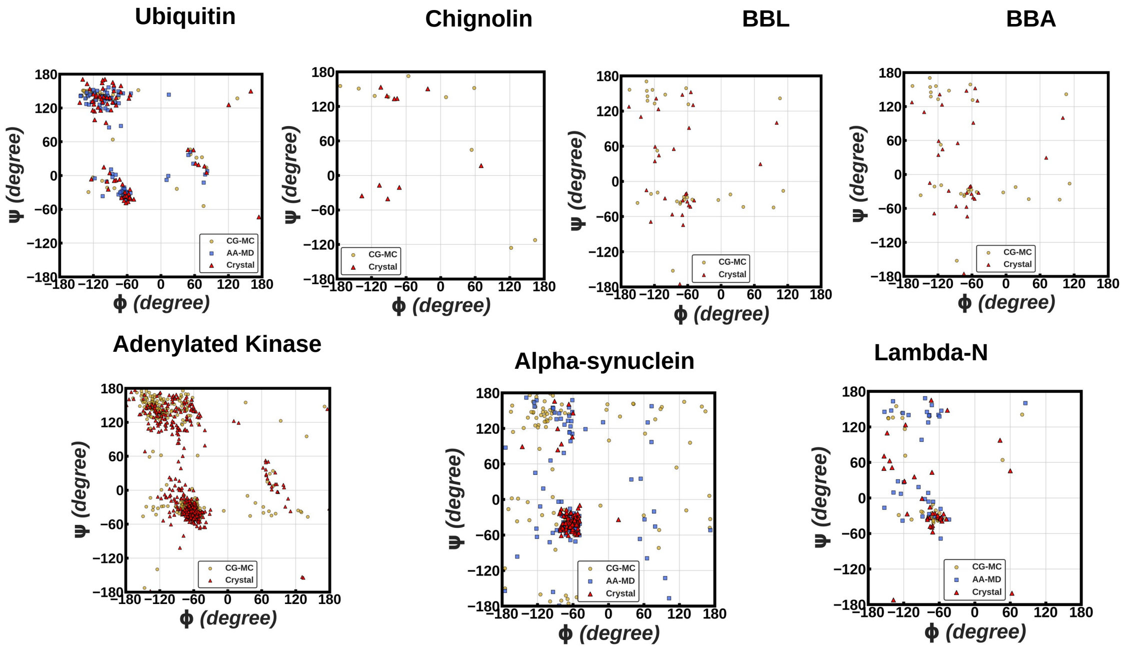


Figure S4: Comparison of Ramachandran (RC) plots for Ubiquitin, Chignolin, BBL, BBA, Adenylated Kinase, $\alpha$-synuclein and $\lambda$N protein obtained from the crystal structure and the final structure from coarse-grained Monte Carlo (CG MC) simulations. Atomsitic data is also added for Ubiquitin, $\alpha$-synuclein and $\lambda$N proteins.

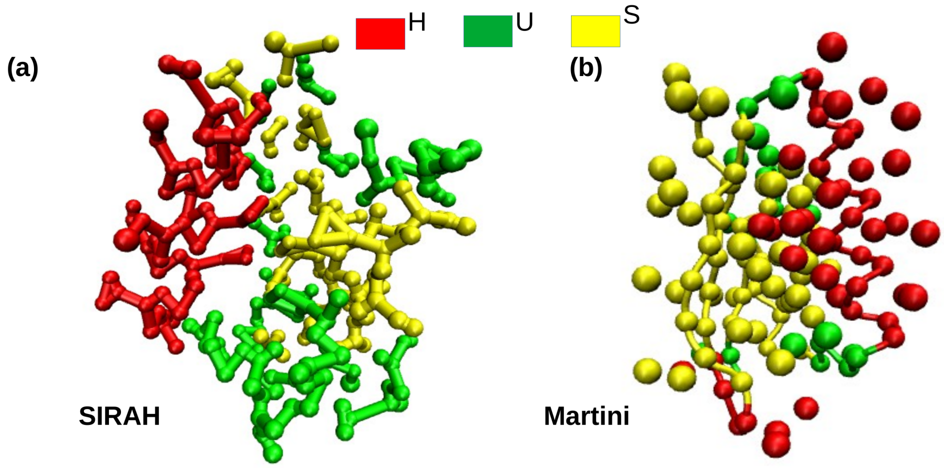


Figure S5: Final GB3 structures after 1 µs simulations using (a) SIRAH and (b) Martini CG force fields. Residues are colored by secondary structure: red for helix, green for unstructured, and yellow for sheet.

Table S1: Classification of GB3 residues as solvophilic or solvophobic in the crystal structure.

| **Res** | **Type** | **Res** | **Type** |
|---|---|---|---|
| MET1 | solvophobic | ALA29 | solvophobic |
| GLN2 | solvophilic | PHE30 | solvophobic |
| TYR3 | solvophobic | LYS31 | solvophilic |
| LYS4 | solvophilic | GLN32 | solvophilic |
| LEU5 | solvophobic | TYR33 | solvophobic |
| VAL6 | solvophobic | ALA34 | solvophobic |
| ILE7 | solvophobic | ASN35 | solvophilic |
| ASN8 | solvophilic | ASP36 | solvophilic |
| GLY9 | solvophobic | ASN37 | solvophilic |
| LYS10 | solvophilic | GLY38 | solvophobic |
| THR11 | solvophilic | VAL39 | solvophobic |
| LEU12 | solvophobic | ASP40 | solvophilic |
| LYS13 | solvophilic | GLY41 | solvophobic |
| GLY14 | solvophobic | VAL42 | solvophobic |
| GLU15 | solvophilic | TRP43 | solvophobic |
| THR16 | solvophilic | THR44 | solvophilic |
| THR17 | solvophilic | TYR45 | solvophobic |
| THR18 | solvophilic | ASP46 | solvophilic |
| LYS19 | solvophilic | ASP47 | solvophilic |
| ALA20 | solvophobic | ALA48 | solvophobic |
| VAL21 | solvophobic | THR49 | solvophilic |
| ASP22 | solvophilic | LYS50 | solvophilic |
| ALA23 | solvophobic | THR51 | solvophilic |
| GLU24 | solvophilic | PHE52 | solvophobic |
| THR25 | solvophilic | THR53 | solvophilic |
| ALA26 | solvophobic | VAL54 | solvophobic |
| GLU27 | solvophilic | THR55 | solvophilic |
| LYS28 | solvophilic | GLU56 | solvophilic |

Table S2: Comparison of secondary structure preferences in GB3 protein. MD denotes trajectory of all atom molecular dynamics simulations and MC denotes conformations based on monte carlo simulations. 'H' corresponds Helix, 'S' corresponds sheet and 'U' corresponds to other than element helix or sheet i.e. loop/coil/turn/bend region of the protein. A bead is assigned the structural element that appears most frequently during the equilibrium trajectory.

| **Res** | **Crystal Structure** | **AA Structure (MD)** | **CG Structure (MC)** | **Res** | **Crystal Structure** | **AA Structure (MD)** | **CG Structure (MC)** |
|---|---|---|---|---|---|---|---|
| MET1 | U | U | U | ALA29 | H | H | H |
| GLN2 | S | S | S | PHE30 | H | H | H |
| TYR3 | S | S | S | LYS31 | H | H | H |
| LYS4 | S | S | S | GLN32 | H | H | H |
| LEU5 | S | U | S | TYR33 | H | H | H |
| VAL6 | S | S | S | ALA34 | H | H | H |
| ILE7 | S | S | S | ASN35 | H | H | H |
| ASN8 | S | U | S | ASP36 | H | H | H |
| GLY9 | U | U | S | ASN37 | H | H | H |
| LYS10 | U | U | U | GLY38 | U | U | U |
| THR11 | U | U | U | VAL39 | U | U | S |
| LEU12 | U | U | S | ASP40 | U | U | S |
| LYS13 | S | U | S | GLY41 | U | U | S |
| GLY14 | S | U | U | VAL42 | S | S | S |
| GLU15 | S | S | S | TRP43 | S | S | S |
| THR16 | S | S | S | THR44 | S | S | S |
| THR17 | S | S | S | TYR45 | S | S | S |
| THR18 | S | S | S | ASP46 | S | S | S |
| LYS19 | S | S | S | ASP47 | U | U | U |
| ALA20 | U | U | S | ALA48 | U | U | U |
| VAL21 | U | U | U | THR49 | U | U | U |
| ASP22 | H | U | S | LYS50 | U | U | U |
| ALA23 | H | H | H | THR51 | S | S | S |
| GLU24 | H | H | H | PHE52 | S | S | S |
| THR25 | H | H | H | THR53 | S | S | S |
| ALA26 | H | H | H | VAL54 | S | S | S |
| GLU27 | H | H | H | THR55 | S | S | U |
| LYS28 | H | H | H | GLU56 | S | U | U |

Table S3: Comparison of secondary structure preferences in GB3 protein (random shift crystal dihedral within a range of $-\pi/10$ to $+\pi/10$). MD denotes trajectory of all atom molecular dynamics simulations and MC denotes conformations based on Monte Carlo simulations. 'H' corresponds Helix, 'S' corresponds sheet and 'U' corresponds to other than element helix or sheet i.e. loop/coil/turn/bend region of the protein. Each bead is assigned the structural element that occurs most frequently over the equilibrium trajectory.

| **Res** | **Crystal Structure** | **AA Structure (MD)** | **CG Structure (MC)** | **Res** | **Crystal Structure** | **AA Structure (MD)** | **CG Structure (MC)** |
|---|---|---|---|---|---|---|---|
| MET1 | U | U | U | ALA29 | H | H | H |
| GLN2 | S | S | S | PHE30 | H | H | H |
| TYR3 | S | S | S | LYS31 | H | H | H |
| LYS4 | S | S | S | GLN32 | H | H | H |
| LEU5 | S | U | U | TYR33 | H | H | H |
| VAL6 | S | S | S | ALA34 | H | H | H |
| ILE7 | S | S | S | ASN35 | H | H | H |
| ASN8 | S | U | S | ASP36 | H | H | H |
| GLY9 | U | U | S | ASN37 | H | H | H |
| LYS10 | U | U | U | GLY38 | U | U | U |
| THR11 | U | U | U | VAL39 | U | U | S |
| LEU12 | U | U | S | ASP40 | U | U | S |
| LYS13 | S | U | S | GLY41 | U | U | S |
| GLY14 | S | U | U | VAL42 | S | S | S |
| GLU15 | S | S | S | TRP43 | S | S | S |
| THR16 | S | S | S | THR44 | S | S | S |
| THR17 | S | S | S | TYR45 | S | S | S |
| THR18 | S | S | S | ASP46 | S | S | S |
| LYS19 | S | S | S | ASP47 | U | U | U |
| ALA20 | U | U | S | ALA48 | U | U | U |
| VAL21 | U | U | U | THR49 | U | U | U |
| ASP22 | H | U | U | LYS50 | U | U | U |
| ALA23 | H | H | H | THR51 | S | S | S |
| GLU24 | H | H | H | PHE52 | S | S | S |
| THR25 | H | H | H | THR53 | S | S | S |
| ALA26 | H | H | H | VAL54 | S | S | S |
| GLU27 | H | H | H | THR55 | S | S | U |
| LYS28 | H | H | H | GLU56 | S | U | U |

Table S4: Comparison of secondary structure preferences in GB3 protein using CHARMM force field parameters for bonded interactions in CG simulations and for all interactions in AA MD simulations. MD denotes trajectory of all atom molecular dynamics simulations and MC denotes conformations based on Monte Carlo simulations. 'H' corresponds Helix, 'S' corresponds sheet and 'U' corresponds to other than element helix or sheet i.e. loop/coil/turn/bend region of the protein. Each bead is assigned its most frequently observed structural element during the equilibrium trajectory.

| **Res** | **Crystal Structure** | **AA Structure (MD)** | **CG Structure (MC)** | **Res** | **Crystal Structure** | **AA Structure (MD)** | **CG Structure (MC)** |
|---|---|---|---|---|---|---|---|
| MET1 | U | U | U | ALA29 | H | H | H |
| GLN2 | S | S | S | PHE30 | H | H | H |
| TYR3 | S | S | S | LYS31 | H | H | H |
| LYS4 | S | S | S | GLN32 | H | H | H |
| LEU5 | S | S | U | TYR33 | H | H | H |
| VAL6 | S | S | S | ALA34 | H | H | H |
| ILE7 | S | S | S | ASN35 | H | H | H |
| ASN8 | S | U | U | ASP36 | H | H | H |
| GLY9 | U | U | U | ASN37 | H | H | H |
| LYS10 | U | U | U | GLY38 | U | U | U |
| THR11 | U | U | U | VAL39 | U | U | S |
| LEU12 | U | U | S | ASP40 | U | U | S |
| LYS13 | S | U | S | GLY41 | U | U | U |
| GLY14 | S | U | U | VAL42 | S | S | S |
| GLU15 | S | S | S | TRP43 | S | S | S |
| THR16 | S | S | S | THR44 | S | S | S |
| THR17 | S | S | S | TYR45 | S | S | U |
| THR18 | S | S | S | ASP46 | S | S | U |
| LYS19 | S | S | S | ASP47 | U | U | U |
| ALA20 | U | U | S | ALA48 | U | U | U |
| VAL21 | U | U | U | THR49 | U | U | U |
| ASP22 | H | U | U | LYS50 | U | U | U |
| ALA23 | H | H | H | THR51 | S | S | U |
| GLU24 | H | H | H | PHE52 | S | U | S |
| THR25 | H | H | H | THR53 | S | S | U |
| ALA26 | H | H | H | VAL54 | S | S | S |
| GLU27 | H | H | H | THR55 | S | S | U |
| LYS28 | H | H | H | GLU56 | S | U | U |

Table S5: Comparison of secondary structure preferences for homeodomain protein. MD denotes trajectory of all atom molecular dynamics simulations and MC denotes conformations based on Monte Carlo simulations. 'H' corresponds Helix, 'S' corresponds sheet and 'U' corresponds to other than element helix or sheet i.e. loop/coil/turn/bend region of the protein. Each bead is assigned its preferred structural element, defined as the one that occurs most frequently over the equilibrium trajectory.

| **Res** | **Crystal Structure** | **AA Structure (MD)** | **CG Structure (MC)** | **Res** | **Crystal Structure** | **AA Structure (MD)** | **CG Structure (MC)** |
|---|---|---|---|---|---|---|---|
| ARG1 | U | U | U | GLY30 | H | H | U |
| GLY2 | U | U | U | LEU31 | H | H | H |
| HIS3 | U | U | U | GLU32 | H | H | H |
| ARG4 | U | H | U | ASN33 | H | H | H |
| PHE5 | U | U | U | LEU34 | H | H | H |
| THR6 | U | U | U | MET35 | H | H | H |
| ALA7 | H | H | H | LYS36 | H | H | H |
| GLU8 | H | H | H | ASN37 | H | H | H |
| ASN9 | H | H | H | THR38 | H | H | H |
| VAL10 | H | H | H | SER39 | U | U | H |
| ARG11 | H | H | H | LEU40 | U | U | U |
| ILE12 | H | H | H | SER41 | U | U | U |
| LEU13 | H | H | H | ARG42 | H | H | H |
| GLU14 | H | H | H | ILE43 | H | H | H |
| SER15 | H | H | H | GLN44 | H | H | H |
| TRP16 | H | H | H | ILE45 | H | H | H |
| PHE17 | H | H | H | LYS46 | H | H | H |
| ALA18 | H | H | H | ASN47 | H | H | H |
| ALA19 | H | H | H | TRP48 | H | H | H |
| ASN20 | H | H | H | VAL49 | H | H | H |
| ILE21 | U | H | H | SER50 | H | H | H |
| ALA22 | U | H | H | ASN51 | H | H | H |
| ASN23 | U | U | U | ARG52 | H | H | H |
| PRO24 | U | U | U | ARG53 | H | U | H |
| TYR25 | U | U | U | ARG54 | H | U | H |
| LEU26 | U | U | U | LYS55 | H | U | H |
| ASP27 | U | U | U | GLU56 | H | U | H |
| THR28 | H | U | H | ALA57 | U | U | H |
| LYS29 | H | U | H | ALA58 | U | U | H |

Table S6: Comparison of secondary structure preferences for ubiquitin protein. MD denotes trajectory of all atom molecular dynamics simulations and MC denotes conformations based on Monte Carlo simulations. 'H' corresponds Helix, 'S' corresponds sheet and 'U' corresponds to other than element helix or sheet i.e. loop/coil/turn/bend region of the protein. Each bead is assigned the structural element that appears most frequently during the equilibrium trajectory.

| **Res** | **Crystal Structure** | **AA Structure (MD)** | **CG Structure (MC)** | **Res** | **Crystal Structure** | **AA Structure (MD)** | **CG Structure (MC)** |
|---|---|---|---|---|---|---|---|
| MET1 | U | U | U | ASP39 | H | H | H |
| GLN2 | S | S | S | GLN40 | H | H | H |
| ILE3 | S | S | S | GLN41 | S | S | S |
| PHE4 | S | S | S | ARG42 | S | S | S |
| VAL5 | S | S | S | LEU43 | S | S | S |
| LYS6 | S | S | S | ILE44 | S | S | S |
| THR7 | U | S | S | PHE45 | S | S | S |
| LEU8 | U | H | H | ALA46 | U | U | U |
| THR9 | U | U | H | GLY47 | U | H | U |
| GLY10 | U | U | U | LYS48 | S | S | S |
| LYS11 | U | S | S | GLN49 | S | S | S |
| THR12 | S | S | S | LEU50 | U | S | S |
| ILE13 | S | S | S | GLN51 | U | S | S |
| THR14 | S | S | S | ASP52 | U | H | H |
| LEU15 | S | S | S | GLY53 | U | H | H |
| GLU16 | S | S | S | ARG54 | U | S | S |
| VAL17 | U | S | S | THR55 | S | S | S |
| GLU18 | U | S | S | LEU56 | H | H | H |
| PRO19 | U | H | H | SER57 | H | H | H |
| SER20 | U | H | H | ASP58 | H | H | H |
| ASP21 | U | S | S | TYR59 | H | H | H |
| THR22 | S | S | S | ASN60 | U | H | H |
| ILE23 | H | H | H | ILE61 | U | S | S |
| GLU24 | H | H | H | GLN62 | U | S | S |
| ASN25 | H | H | H | LYS63 | U | S | S |
| VAL26 | H | H | H | GLU64 | U | H | U |
| LYS27 | H | H | H | SER65 | U | S | S |
| ALA28 | H | H | H | THR66 | S | S | S |
| LYS29 | H | H | H | LEU67 | S | S | S |
| ILE30 | H | H | H | HIS68 | S | S | S |
| GLN31 | H | H | H | LEU69 | S | S | S |
| ASP32 | H | H | H | VAL70 | S | S | S |
| LYS33 | H | H | H | LEU71 | S | S | S |
| GLU34 | H | H | H | ARG72 | U | U | S |
| GLY35 | U | H | H | LEU73 | U | U | S |
| ILE36 | U | S | S | ARG74 | U | U | S |
| PRO37 | U | S | S | GLY75 | U | U | U |
| PRO38 | H | H | H | GLY76 | U | U | U |

Table S7. Comparison of secondary-structure preferences in the Chignolin protein. MC denotes conformations generated from Monte Carlo simulations. 'H' represents helix, 'S' represents $\beta$-sheet and 'U' represents regions other than helix or sheet, including loop, coil, turn and bend structures. A bead is assigned the secondary-structure element that appears most frequently during the equilibrium trajectory.

| **Res** | **Crystal Structure** | **CG Structure (MC)** | **Res** | **Crystal Structure** | **CG Structure (MC)** |
|---|---|---|---|---|---|
| TYR1 | U | U | THR6 | U | U |
| TYR2 | S | S | GLY7 | U | U |
| ASP3 | U | U | THR8 | U | U |
| PRO4 | U | U | TRP9 | S | U |
| GLU5 | U | U | TYR10 | U | U |

Table S8. Comparison of secondary-structure preferences in the BBL protein. MC denotes conformations generated from Monte Carlo simulations. 'H' represents helix, 'S' represents $\beta$-sheet and 'U' represents regions other than helix or sheet, including loop, coil, turn and bend structures. A bead is assigned the secondary-structure element that appears most frequently during the equilibrium trajectory.

| **Res** | **Crystal Structure** | **CG Structure (MC)** | **Res** | **Crystal Structure** | **CG Structure (MC)** |
|---|---|---|---|---|---|
| GLY1 | U | U | ALA25 | H | U |
| SER2 | U | U | ILE26 | U | U |
| GLN3 | U | U | LYS27 | U | U |
| ASN4 | U | U | GLY28 | U | U |
| ASN5 | U | U | THR29 | U | U |
| PHE6 | U | U | GLY30 | U | U |
| ALA7 | U | U | VAL31 | U | U |
| LEU8 | U | U | GLY32 | U | U |
| SER9 | U | H | GLY33 | U | U |
| PRO10 | H | H | ARG34 | U | U |
| ALA11 | H | H | LEU35 | U | U |
| ILE12 | H | H | THR36 | U | U |
| ARG13 | H | H | ARG37 | H | H |
| ARG14 | H | H | GLU38 | H | H |
| LEU15 | H | H | ASP39 | H | U |
| LEU16 | H | H | VAL40 | H | H |
| ALA17 | H | U | GLU41 | H | U |
| GLU18 | H | H | LYS42 | H | H |
| TRP19 | H | U | HIS43 | H | U |
| ASN20 | U | U | LEU44 | H | U |
| LEU21 | U | U | ALA45 | H | U |
| ASP22 | U | U | LYS46 | H | U |
| ALA23 | H | H | ALA47 | U | U |
| SER24 | H | U | - | - | - |

Table S9. Comparison of secondary-structure preferences in the BBA protein. MC denotes conformations generated from Monte Carlo simulations. 'H' represents helix, 'S' represents $\beta$-sheet and 'U' represents regions other than helix or sheet, including loop, coil, turn, and bend structures. A bead is assigned the secondary-structure element that appears most frequently during the equilibrium trajectory.

| **Res** | **Crystal Structure** | **CG Structure (MC)** | **Res** | **Crystal Structure** | **CG Structure (MC)** |
|---|---|---|---|---|---|
| GLU1 | U | U | GLU15 | H | U |
| GLN2 | U | S | LYS16 | H | U |
| TYR3 | U | U | GLU17 | H | H |
| THR4 | U | U | LEU18 | H | H |
| ALA5 | U | U | ARG19 | H | H |
| LYS6 | S | S | ASP20 | U | U |
| TYR7 | U | U | PHE21 | H | H |
| LYS8 | U | U | ILE22 | H | U |
| GLY9 | U | U | GLU23 | H | U |
| ARG10 | S | S | LYS24 | H | H |
| THR11 | S | S | PHE25 | H | U |
| PHE12 | S | U | LYS26 | U | U |
| ARG13 | U | U | GLY27 | U | U |
| ASN14 | U | U | ARG28 | U | U |

Table S10 (a). Comparison of secondary structure elements of the di-ubiquitin protein on a per-residue basis. MD refers to the all-atom molecular dynamics (AA MD) trajectory, and MC represents conformations obtained from coarse-grained Monte Carlo (CG MC) simulations. 'H' denotes helix, 'S' denotes sheet, and 'U' represents unstructured regions (loop/coil/turn/bend). Each bead is assigned the structural element that appears most frequently during the equilibrium trajectory.

| **Res** | **Initial Structure** | **AA Structure (MD)** | **CG Structure (MC)** | **Res** | **Initial Structure** | **AA Structure (MD)** | **CG Structure (MC)** |
|---|---|---|---|---|---|---|---|
| MET1 | U | U | U | MET1 | U | U | U |
| GLN2 | S | S | S | GLN2 | S | S | U |
| ILE3 | S | S | S | ILE3 | S | S | S |
| PHE4 | S | S | S | PHE4 | S | S | S |
| VAL5 | S | S | S | VAL5 | S | S | S |
| LYS6 | S | S | U | LYS6 | S | S | U |
| THR7 | U | S | U | THR7 | U | S | S |
| LEU8 | U | H | H | LEU8 | U | H | H |
| THR9 | U | H | H | THR9 | U | U | U |
| GLY10 | U | U | U | GLY10 | U | U | U |
| LYS11 | U | U | U | LYS11 | U | S | S |
| THR12 | S | S | S | THR12 | S | S | S |
| ILE13 | S | S | S | ILE13 | S | S | S |
| THR14 | S | S | S | THR14 | S | S | S |
| LEU15 | S | S | S | LEU15 | S | S | S |
| GLU16 | S | S | S | GLU16 | S | S | S |
| VAL17 | U | U | S | VAL17 | U | S | S |
| GLU18 | U | S | S | GLU18 | U | S | S |
| PRO19 | U | H | H | PRO19 | U | H | U |
| SER20 | U | U | H | SER20 | U | U | H |
| ASP21 | U | S | U | ASP21 | U | S | U |
| THR22 | S | S | S | THR22 | S | S | S |
| ILE23 | H | H | H | ILE23 | H | H | U |
| GLU24 | H | H | H | GLU24 | H | H | H |
| ASN25 | H | H | H | ASN25 | H | H | H |
| VAL26 | H | H | H | VAL26 | H | H | H |
| LYS27 | H | H | H | LYS27 | H | H | H |
| ALA28 | H | H | H | ALA28 | H | H | H |
| LYS29 | H | H | H | LYS29 | H | H | H |
| ILE30 | H | H | H | ILE30 | H | H | H |
| GLN31 | H | H | H | GLN31 | H | H | H |
| ASP32 | H | H | H | ASP32 | H | H | H |
| LYS33 | H | H | H | LYS33 | H | U | H |
| GLU34 | H | U | U | GLU34 | H | U | U |
| GLY35 | U | U | U | GLY35 | U | U | U |
| ILE36 | U | U | U | ILE36 | U | S | U |
| PRO37 | U | U | U | PRO37 | U | U | S |
| PRO38 | U | H | H | PRO38 | H | H | U |
| ASP39 | U | H | H | ASP39 | H | H | H |
| GLN40 | U | U | U | GLN40 | H | U | H |

Table S10 (b). Comparison of secondary structure elements of the di-ubiquitin protein on a per-residue basis. MD refers to the all-atom molecular dynamics (AA MD) trajectory, and MC represents conformations obtained from coarse-grained Monte Carlo (CG MC) simulations. 'H' denotes helix, 'S' denotes sheet, and 'U' represents unstructured regions (loop/coil/turn/bend). Each bead is assigned the structural element that occurs most frequently over the equilibrium trajectory.

| **Res** | **Initial Structure** | **AA Structure (MD)** | **CG Structure (MC)** | **Res** | **Initial Structure** | **AA Structure (MD)** | **CG Structure (MC)** |
|---|---|---|---|---|---|---|---|
| GLN41 | S | S | S | GLN41 | S | S | H |
| ARG42 | S | S | S | ARG42 | S | S | S |
| LEU43 | S | S | S | LEU43 | S | S | S |
| ILE44 | S | S | S | ILE44 | S | S | S |
| PHE45 | S | U | U | PHE45 | S | U | S |
| ALA46 | U | U | U | ALA46 | U | U | S |
| GLY47 | U | U | U | GLY47 | U | U | U |
| ARG48 | S | S | U | ARG48 | S | S | U |
| GLN49 | S | S | U | GLN49 | S | U | S |
| LEU50 | U | S | S | LEU50 | U | S | U |
| GLU51 | U | S | S | GLU51 | U | S | S |
| ASP52 | U | U | H | ASP52 | U | H | U |
| GLY53 | U | U | H | GLY53 | U | U | U |
| ARG54 | U | S | S | ARG54 | U | S | U |
| THR55 | S | U | U | THR55 | S | S | S |
| LEU56 | H | H | H | LEU56 | H | H | U |
| SER57 | H | H | H | SER57 | H | H | H |
| ASP58 | H | H | H | ASP58 | H | H | H |
| TYR59 | H | U | U | TYR59 | H | U | H |
| ASN60 | U | U | U | ASN60 | U | U | H |
| ILE61 | U | U | U | ILE61 | U | S | U |
| GLN62 | U | S | S | GLN62 | U | S | U |
| LYS63 | U | U | S | LYS63 | U | U | S |
| GLU64 | U | U | U | GLU64 | U | U | U |
| SER65 | U | S | U | SER65 | U | S | U |
| THR66 | S | S | U | THR66 | S | S | S |
| LEU67 | S | S | S | LEU67 | S | S | S |
| HIS68 | S | S | S | HIS68 | S | S | S |
| LEU69 | S | S | S | LEU69 | S | S | S |
| VAL70 | S | S | S | VAL70 | S | S | S |
| LEU71 | S | S | S | LEU71 | S | S | S |
| ARG72 | U | U | S | ARG72 | U | S | S |
| LEU73 | U | U | U | LEU73 | U | U | S |
| | | | | ARG74 | U | U | U |
| | | | | GLY75 | U | U | U |

Table S11 (a). Comparison of secondary structure elements of the $\alpha$-synuclein protein on a per-residue basis. MD refers to the all-atom molecular dynamics (AA MD) trajectory, and MC represents conformations obtained from coarse-grained Monte Carlo (CG MC) simulations. 'H' denotes helix, 'S' denotes sheet, and 'U' represents unstructured regions (loop/coil/turn/bend).

| **Res** | **Initial Structure** | **AA Structure (MD)** | **CG Structure (MC)** | **Res** | **Initial Structure** | **AA Structure (MD)** | **CG Structure (MC)** |
|---|---|---|---|---|---|---|---|
| MET1 | U | U | U | VAL37 | H | U | U |
| ASP2 | U | U | U | LEU38 | U | U | U |
| VAL3 | H | U | U | TYR39 | U | U | U |
| PHE4 | H | U | U | VAL40 | U | U | U |
| MET5 | H | U | U | GLY41 | U | U | U |
| LYS6 | H | U | U | SER42 | U | U | U |
| GLY7 | H | U | U | LYS43 | U | U | U |
| LEU8 | H | U | U | THR44 | U | U | U |
| SER9 | H | U | U | LYS45 | H | U | U |
| LYS10 | H | U | U | GLU46 | H | U | U |
| ALA11 | H | U | U | GLY47 | H | U | U |
| LYS12 | H | U | U | VAL48 | H | H | U |
| GLU13 | H | U | U | VAL49 | H | H | U |
| GLY14 | H | U | U | HIS50 | H | H | U |
| VAL15 | H | U | U | GLY51 | H | H | U |
| VAL16 | H | H | U | VAL52 | H | H | U |
| ALA17 | H | H | U | ALA53 | H | H | U |
| ALA18 | H | H | U | THR54 | H | U | U |
| ALA19 | H | H | U | VAL55 | H | U | U |
| GLU20 | H | H | U | ALA56 | H | U | U |
| LYS21 | H | H | U | GLU57 | H | H | U |
| THR22 | H | H | U | LYS58 | H | U | U |
| LYS23 | H | H | U | THR59 | H | U | U |
| GLN24 | H | H | U | LYS60 | H | U | U |
| GLY25 | H | H | U | GLU61 | H | U | U |
| VAL26 | H | H | U | GLN62 | H | U | U |
| ALA27 | H | H | U | VAL63 | H | U | U |
| GLU28 | H | H | U | THR64 | H | U | U |
| ALA29 | H | H | U | ASN65 | H | U | U |
| ALA30 | H | U | U | VAL66 | H | U | U |
| GLY31 | H | U | U | GLY67 | H | U | U |
| LYS32 | H | U | U | GLY68 | H | U | U |
| THR33 | H | U | U | ALA69 | H | H | U |
| LYS34 | H | U | U | VAL70 | H | U | U |
| GLU35 | H | U | U | VAL71 | H | U | U |
| GLY36 | H | U | U | THR72 | H | U | U |
| GLY73 | H | U | U | ALA85 | H | U | U |
| VAL74 | H | U | U | GLY86 | H | U | U |

Table S11 (b). Comparison of secondary structure elements of the $\alpha$-synuclein protein on a per-residue basis. MD refers to the all-atom molecular dynamics (AA MD) trajectory, and MC represents conformations obtained from coarse-grained Monte Carlo (CG MC) simulations. 'H' denotes helix, 'S' denotes sheet, and 'U' represents unstructured regions (loop/coil/turn/bend). Each bead is assigned the structural element that occurs most frequently over the equilibrium trajectory.

| **Res** | **Initial Structure** | **AA Structure (MD)** | **CG Structure (MC)** | **Res** | **Initial Structure** | **AA Structure (MD)** | **CG Structure (MC)** |
|---|---|---|---|---|---|---|---|
| THR75 | H | U | U | SER87 | H | U | U |
| ALA76 | H | U | U | ILE88 | H | H | U |
| VAL77 | H | U | U | ALA89 | H | H | U |
| ALA78 | H | U | U | ALA90 | H | H | U |
| GLN79 | H | H | U | ALA91 | H | H | U |
| LYS80 | H | H | U | THR92 | H | U | U |
| THR81 | H | H | U | GLY93 | U | U | U |
| VAL82 | H | H | U | PHE94 | U | U | U |
| GLU83 | H | U | U | VAL95 | U | U | H |
| GLY84 | H | U | U | | | | |

Table S12. Comparison of secondary structure elements of the λN protein on a per-residue basis. MD refers to the all-atom molecular dynamics (AA MD) trajectory, and MC represents conformations obtained from coarse-grained Monte Carlo (CG MC) simulations. 'H' denotes helix, 'S' denotes sheet, and 'U' represents unstructured regions (loop/coil/turn/bend). As λN is an IDP, the percentage (%) of each conformation per residue is reported. All values within brackets indicate percentages. Each bead is assigned its most frequently observed structural element during the equilibrium trajectory.

| **Res** | **Initial Structure** | **AA Structure (MD)** | **CG Structure (MC)** | **Res** | **Initial Structure** | **AA Structure (MD)** | **CG Structure (MC)** |
|---|---|---|---|---|---|---|---|
| ASP1 | U | U | U | ALA19 | H | U | U |
| ALA2 | H | U | U | ALA20 | H | U | U |
| GLN3 | H | U | U | ASN21 | U | U | U |
| THR4 | H | U | U | PRO22 | H | U | U |
| ARG5 | H | U | U | LEU23 | H | U | U |
| ARG6 | H | U | U | LEU24 | H | U | U |
| ARG7 | H | U | U | VAL25 | U | U | U |
| GLU8 | H | U | U | GLY26 | U | U | U |
| ARG9 | H | U | U | VAL27 | U | U | U |
| ARG10 | H | U | U | SER28 | U | U | U |
| ALA11 | H | U | U | ALA29 | U | U | U |
| GLU12 | H | U | U | LYS30 | U | U | U |
| LYS13 | H | U | U | PRO31 | U | U | U |
| GLN14 | H | U | U | VAL32 | U | U | U |
| ALA15 | H | U | U | ASN33 | U | U | U |
| GLN16 | H | U | U | ARG34 | U | U | U |
| TRP17 | H | U | U | PRO35 | U | U | U |
| LYS18 | H | U | U | | | | |

(3) Van Der Spoel, D.; Lindahl, E.; Hess, B.; Groenhof, G.; Mark, A. E.; Berendsen, H. J.

GROMACS: fast, flexible, and free. *Journal of computational chemistry* **2005**, *26*, 1701–1718.
